# Lower statistical support with larger datasets: insights from the Ochrophyta radiation

**DOI:** 10.1101/2021.01.14.426536

**Authors:** Arnaud Di Franco, Denis Baurain, Gernot Glöckner, Michael Melkonian, Hervé Philippe

**Author notes:** deceased (Gernot Glöckner).

## Abstract

It is commonly assumed that increasing the number of characters has the potential to resolving radiations. We studied photosynthetic stramenopiles (Ochrophyta) using alignments of heterogeneous size and origin (6,762 sites for mitochondrion, 21,692 sites for plastid and 209,105 sites for nucleus). While statistical support for the relationships between the six major Ochrophyta lineages increases when comparing the mitochondrion and plastid trees, it decreases in the nuclear tree. Statistical support is not simply related to the dataset size but also to the quantity of phylogenetic signal available at each position and our ability to extract it. Here, we show that proper signal extraction is difficult to attain, as demonstrated by conflicting results obtained when varying taxon sampling. Even though the use of a better fitting model improved signal extraction and reduced the observed conflicts, the plastid dataset provided higher statistical support for the ochrophyte radiation than the larger nucleus dataset. We propose that the higher support observed in the plastid tree is due to an acceleration of the evolutionary rate in one short deep internal branch, implying that more phylogenetic signal per position is available to resolve the Ochrophyta radiation in the plastid than in the nuclear dataset. Our work therefore suggests that, in order to resolve radiations, beyond the obvious use of datasets with more positions, we need to continue developing models of sequence evolution that better extract the phylogenetic signal and design methods to search for genes/characters that contain more signal specifically for short internal branches.

## Introduction

One of the last major challenges of phylogenetics is to resolve ancient radiations. They combine three main difficulties encountered during phylogenetic inference. First, they are by definition characterized by short internal branches, meaning a scarce genuine (historical) phylogenetic signal, which resides in the rare substitutions accumulated during a short amount of time (i.e., very few synapomorphies). Second, their ancient nature makes the sites subject to substitutional saturation (multiple substitutions at the same site). As a result, an ancient synapomorphy can easily be masked by subsequent substitutions, leading the tree reconstruction method to possibly interpret it as a convergence. The misinterpretation of site history, if not random (i.e., biased), creates a non-phylogenetic signal that conflicts with the genuine phylogenetic signal. Third, all loci possibly do not share the same evolutionary history, because of incomplete lineage sorting (ILS) or even more problematically, hybridisation (Maddison1997). To sum up, the phylogenetic signal left behind by ancient radiations is both scarce and difficult to extract (Whitfield and Lockhart 2007). Because increasing the size of the dataset is a necessary condition to resolve such radiations, phylogenomics, the use of large datasets, initially generated great hope (Gee 2003). However, the failure of phylogenomics to resolve most of those complex cases suggests that more data may not be sufficient (see Philippe et al. 2011).

This failure might be due to limitations of existing tree reconstruction methods. Let us consider a dataset D containing n sites. To simplify our initial reasoning, let us assume first that all loci share the same history. The genuine phylogenetic signal (PS) contained in it for a given branch B is:

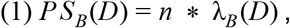

with λ_*B*_ (*D*) being the expected number of substitutions per site in branch B for dataset D. Yet, PS_B_(D), the total number of synapomorphies of the dataset supporting this branch, is actually unknown, as we only have access to the apparent phylogenetic signal (PS’) (Baurain and Philippe 2010) inferred by a specific method m:

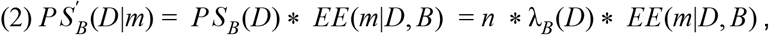

with EE(m |*D*, *B*) being the extraction efficiency of method m for branch B and dataset D. This efficiency depends not only on the properties of the radiation (e.g., age) and the dataset (e.g., global rate of evolution or taxon sampling), but also ultimately on how correctly the model of evolution infers the substitution history at each position. Limitations of tree reconstruction methods generally lead to values of EE(m |*D*, *B*) smaller than one. Conceptually, EE(m |*D*, *B*) might even be negative when a bias favors an alternative branching order (e.g., a long branch attraction between two unrelated fast-evolving taxa) and larger than one when such a bias favors the correct solution (e.g., a long branch attraction between two fast-evolving taxa that are really sisters). In cases where all loci do not share the same history, the extraction efficiency might be further reduced by other model violations, for example when using a concatenated model in the presence of ILS. The failure to resolve most radiations (i.e., PS’(D|m) not significantly different from 0, as often revealed by a low statistical support) is due to the fact that n*λ_B_(D) and/or EE(m |*D*, *B*) are too small. In phylogenomics, n tends to be maximal and improvements on n*λ_B_(D) become asymptotically smaller. This is especially true when considering the finite collection of orthologous sequences relevant to the issue at hand. While alternative approaches based on other types of characters may exist (e.g., retrotransposon insertions or intron positions), they will not be considered here. The hope of supermatrix-based phylogenomics to resolve radiations mainly hinges on improving the extraction efficiency of the tree inference methods.

Since phylogenetic inference should be viewed as a statistical problem (Felsenstein 1983), it requires the formalization of an explicit model. The extraction efficiency of probabilistic tree reconstruction methods ultimately depends on the validity of model assumptions (Simion et al. 2020). In other words, model violations decrease extraction efficiency. Two main types of violations exist: (1) violations of the model of sequence evolution and (2) violations of the model of gene evolution. The first type of violation is unavoidable, as we fail to fully apprehend the complexity of sequence evolution, and it affects all tree reconstruction methods. The second violation is due to the fact that, because of ILS, gene duplication/loss, horizontal gene transfer (HGT) or hybridisation (*i.e.*, gene flux between closely related species), single-gene trees might differ from the species tree (Maddison 1997), which is not taken into account by the concatenation approach. In theory, gene duplication and HGT are not present, given that only orthologs should be included in the supermatrix. However, ILS can affect orthologs and is expected to be all the more frequent in phylogenies with short durations between speciation events. The effect of these model violations can be studied through the comparison of trees inferred by models fitting data to a different degree or the variation of taxon sampling.

To study the impact of extraction efficiency on the power of phylogenomics to resolve ancient radiations, we chose the diversification of Ochrophyta (i.e., photosynthetic Stramenopiles). Stramenopiles (also known as heterokonts) is a eukaryotic clade composed mostly (but not only, e.g., kelps) of unicellular species, and is closely related to Alveolata and Rhizaria, the three clades forming the supergroup SAR (Burki et al. 2007). Inside Stramenopiles, Ochrophyta is a monophyletic group of photosynthetic organisms that appeared around 500 MYa (Brown and Sorhannus 2010). The diversity of this clade is large, ranging from the picoplanktonic *Nannochloropsis* (Eustigmatophyceae) to ecologically important diatoms (Bacillariophyta) and multicellular brown algae (Phaeophyceae). As photosynthetic eukaryotes, they harbor three genomic compartments, a nucleus (nu), a mitochondrion (mt) and a plastid (cp), the latter inherited from a red algal endosymbiont (Archibald 2015).

The diversification of the major ochrophyte lineages seems to have occurred relatively rapidly, as demonstrated by the corresponding short internal branches (Yang et al. 2012; Derelle et al. 2016). The apparent phylogenetic signal for these branches (B) is expected to vary across compartments. First, the three genomes have a quite different size (n_mt_ < n_cp_ ≪ n_nu_), suggesting that *PS*’_B_(*nu*|*m*) ≫ *PS*’_B_(*cp*|*m*) > *PS*’_B_(*mt*|*m*). Second, they have evolved under very different mutation/selection pressures (e.g., different G+C content, presence of recombination in the nucleus but likely not in the organelles), leading to different mean substitution rates (λ_B_(mt) ≠ λ_B_(cp) ≠ λ_B_(nu)) and different extraction efficiencies (EE(m|mt,B) ≠ EE(m|cp,B) ≠ EE(m|nu,B)), due to differences in types and levels of model violations. Although the values of λ_B_(D)*EE(m|D,B) vary across the three genomes, it is difficult to predict whether these variations are major. Moreover, while complex red algae are now thought to have repeatedly exchanged plastids laterally, there is no evidence that it was the case within Ochrophyta (Dorrel et al.et al. 2017; Sibbald and Archibald 2020). Therefore, the latter constitute an interesting case study to evaluate the relative importance of n and λ_B_(D)*EE(m|D,B) in our ability to resolve ancient radiations. In particular, it might allow us to determine whether one should increase n or rather work on improving EE(m|D,B).

For this study, we sequenced mitochondrial and plastid genomes from five ochrophyte species belonging to Chrysophyceae, Dictyochophyceae, Pinguiophyceae and Synurophyceae. From these new data, we built three supermatrices, one for each genomic compartment, all representing most of the major ochrophyte clades. Each dataset was carefully constructed, so as to maximize matrix size and completeness, while minimizing erroneous inclusion of non-orthologous genes, contaminated sequences and sequencing errors. With these three largest stramenopiles supermatrices to date, we studied how extraction efficiency affects phylogenetic inference. We first observed incongruent topologies between the three genomes for deep ochrophyte relationships, along with surprisingly lower bootstrap support for the largest dataset when using the conventional model LG4X (Le et al. 2012). We then studied the impact of model violations by varying taxon sampling and by using an alternative model of sequence evolution. Finally, we explored ways to resolve the deep ochrophyte radiation.

## Results & Discussion

### Recovery of the major ochrophyte clades

We carefully assembled three supermatrices from mitochondrial, plastid and nuclear genome sequences, containing 6,672, 21,692 and 209,105 amino acid positions, respectively (Table 1). They included species from eleven major clades of Ochrophyta: Bacillariophyta, Bolidophyceae, Chrysophyceae, Dictyochophyceae, Eustigmatophyceae, Pelagophyceae, Phaeophyceae, Pinguiophyceae, Raphidophyceae, Synurophyceae and Xanthophyceae. The nuclear dataset also included *Synchroma pusilla*, a species of Synchromophyceae, but some clades (Chrysomeridoephyceae, Phaeothamniophyceae, Schizocladiophyceae, Aurearenophyceae, Phaeosacciophyceae and Chrysoparadoxophyceae) were absent. Mitochondrial, plastid and nuclear phylogenies inferred using the LG4X model (Fig. 1A-C) retrieved the monophyly of all major clades with maximal bootstrap support (BS) except Chrysophyceae (see Supplementary Figures 1-3). Chrysophyceae came out as a monophyletic group in the mitochondrion and plastid datasets (BS=56% and 88%, respectively), but were represented by two species only. In the nuclear phylogeny, which includes 12 chrysophycean species, they were paraphyletic, with Synurophyceae nested within Chrysophyceae, in agreement with previous studies (Yang et al. 2012). Monophyly of Synurophyceae+Chrysophyceae (SC clade) was always maximally supported. In the nuclear dataset, their grouping with Synchromophyceae (SSC clade) was highly supported, as in Yang et al. (2012) and Derelle et al. (2016). This SSC clade was also recovered in a plastid phylogeny built with partial *Synchroma* sequences obtained from RNAseq data (Keeling et al. 2014) (data not shown). Consequently, in the following, we consider the SSC clade as one of the ten major ochrophyte clades contained in our analyses. The PX clade (Phaeophyceae and Xanthophyceae) (Kai et al. 2008) was recovered using all three datasets, as well as their sister relationship with Raphidophyceae (PXR clade) (Graf et al. 2020), but with limited support in the mitochondrial dataset (BS=92% and 71%, respectively). Two other previously reported relationships (Yang et al. 2012; Derelle et al. 2016; Han et al. 2018) were highly supported – monophyly of Pelagophyceae and Dictyochophyceae (PD clade) and monophyly of Bolidophyceae and Bacillariophyta (BB clade) – but again mitochondrial support was not maximal for the PD clade (BS=85%). Inside Bacillariophyta, in our nuclear tree, Coscinodiscophyceae were paraphyletic at the base, followed by a monophyletic group composed of Mediophyceae and Bacillariophyceae, as in Parks et al. (2018). Overall, our results were thus in excellent agreement with existing knowledge.

**Table 1.** Dataset composition summary

| Genomic compartment | number of species | number of amino acid positions | number of genes | missing data (%) |
| --- | --- | --- | --- | --- |
| mitochondrion | 64 | 6,762 | 32 | 5.84 |
| plastid | 63 | 21,692 | 99 | 4.12 |
| nucleus | 124 | 209,105 | 797 | 25.56 |

**Figure 1.**
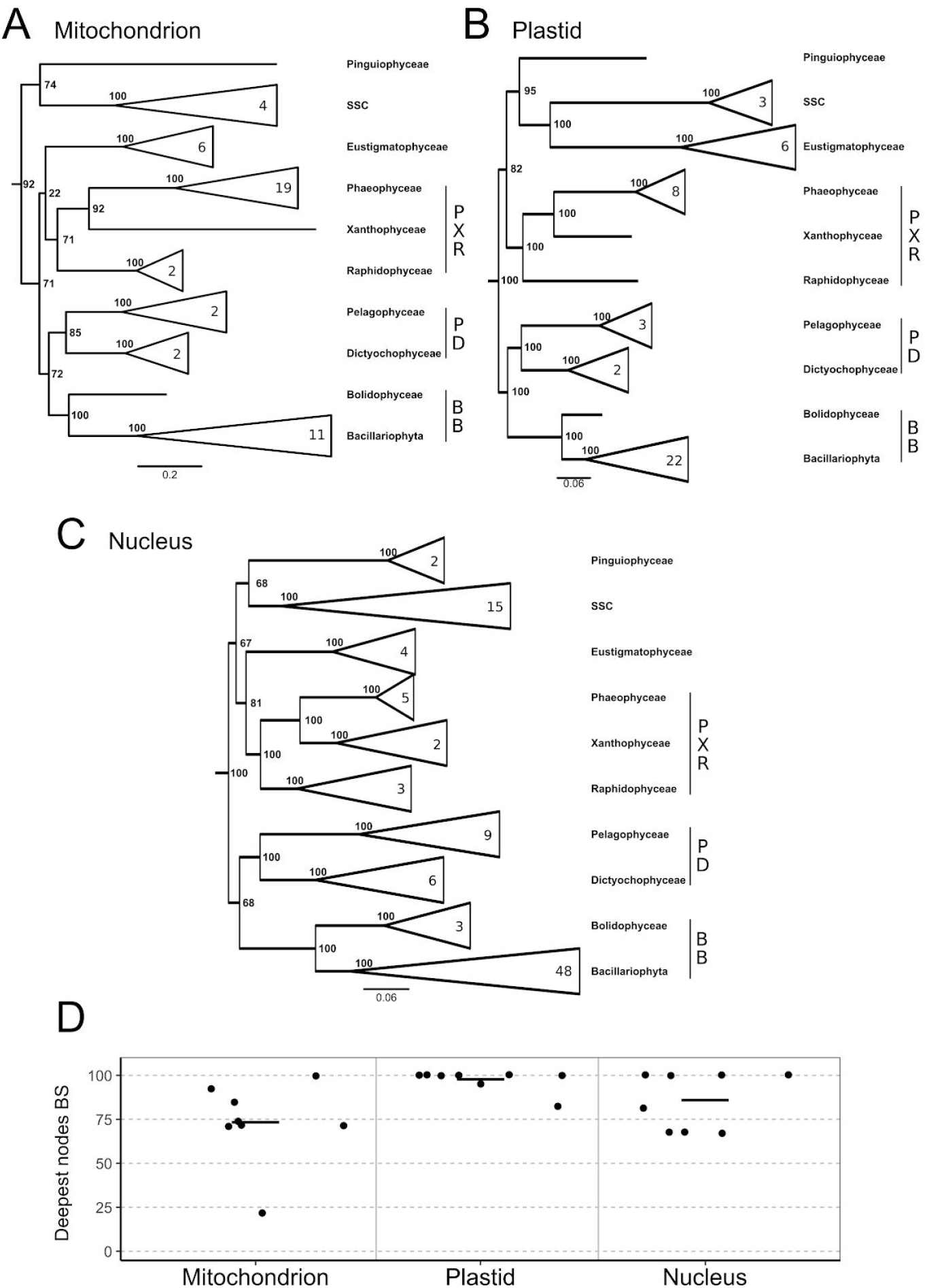
Collapsed trees inferred under the LG4X model from the three different compartments. Statistical support was computed via 100 fast bootstrap replicates in RAxML. The ten major clades of Ochrophyta were collapsed when more than two species were present. SSC stands for the monophyly of Synchromophyceae, Synurophyceae and Chrysophyceae, which is only represented by species of Synurophyceae and Chrysophyceae clades in the mitochondrial and plastid datasets A. mitochondrial dataset - outgroup of 15 species not shown; B. plastid dataset - outgroup of 15 species not shown; C. nuclear dataset - outgroup of 27 species not shown; D. Bootstrap support for eight internal nodes from the three datasets; average values are indicated by a line.

In sharp contrast with the concatenated approach, single-gene phylogenies could only recover the monophyly of the seven well-established, clades (Ochrophyta, BB, Eustigmatophyceae, PD, Pinguiophyceae, PXR and SSC). Whatever the model of sequence evolution used (LG+G, LG4X or CAT+G), their monophylies were found on average ~46% across the 797 nuclear genes, and only 15, 15 and 17 genes recovered all seven clades, respectively. This result is expected given the age of the ochrophyte radiation (~500 MYa) and the limited size of the genes (~260 positions), which favor systematic and stochastic errors respectively. However, this prevents us from studying the impact of ILS on the resolution of Ochrophyta radiation. On one hand, coalescent-based methods that jointly infer gene and species trees (e.g., *BEAST) (Heled and Drummond 2010) are still not accessible to phylogenomics because of their computational burden. On the other hand, the proxies to this joint inference (e.g., ASTRAL) (Mirarab and Warnow 2015; Zhang et al. 2018) are too sensitive to single-gene tree estimation errors to be considered accurate (Gatesy and Springer 2014). Consequently, we cannot study the impact of ILS on the extraction efficiency in the case of the ancient ochrophyte radiation and will focus on the effect of violations of the model of sequence evolution on concatenated data.

### Incongruence between compartments for deep ochrophyte relationships

Although the monophyly of the ten major clades was consistently recovered across the three datasets, the basal phylogeny of Ochrophyta showed incongruent relationships depending on which dataset was used. While the plastid tree strongly grouped Eustigmatophyceae with the SSC clade (BS=100), nuclear and mitochondrial trees separated them, and instead supported the grouping of Pinguiophyceae with SSC on one side (BS=68% and 74%, respectively) and the grouping of Eustigmatophyceae with PXR on the other side (BS=81% and 22%, respectively). Our plastid phylogeny (Fig. 1B) was in agreement with the work of Yang et al. (2012), which was based on nuclear SSU rRNA and four plastid-encoded genes, suggesting that their inferences were dominated by the plastid signal. Comparison with more recently published plastid (Ševčíková et al. 2015), mitochondrial (Ševčíková et al. 2016) and nuclear trees (Derelle et al. 2016) was more difficult, as those datasets lacked some major clades: Bolidophyceae, Dictyochophyceae and Pinguiophyceae (Ševčíková et al. 2015); Bolidophyceae, Dictyochophyceae, Pinguiophyceae and Xanthophyceae (Ševčíková et al. 2016); Eustigmatophyceae and Pinguiophyceae (Derelle et al. 2016). Still, the plastid tree of Ševčíková et al. (2015) was congruent with our plastid tree. However, the mitochondrial tree of Ševčíková et al. (2016) did not recover the monophyly of the PXR clade, contrary to our mitochondrial tree (Fig. 1A), but both trees recovered a basal position for Chrysophyceae. In our nuclear tree (Fig. 1C), we observed the dichotomy between Diatomista (BB+PD) and Chrysista (PXR+SSC), first proposed in Derelle et al. (2016). Overall, although phylogenies based on the three genomic compartments yielded incongruent deep ochrophyte relationships (Fig. 1A-C), they were each in good agreement with previously published trees based on the same compartment (Yang et al. 2012; Ševčíková et al. 2015; Derelle et al. 2016; Ševčíková et al. 2016).

Unexpectedly, statistical support for the eight deep nodes that connect the ten major lineages, displayed a surprising pattern with respect to the number of positions (Fig. 1D). The average BS for these eight nodes increased from 73% in the mitochondrion tree (6,762 positions) to 97% in the plastid tree (21,692 positions), disregarding the fact that these two trees differed for basal relationships. With ~3 times more positions than the mitochondrial dataset, the plastid dataset thus confirmed the expectation that the apparent phylogenetic signal (as measured by bootstrap support) increases with dataset size. In sharp contrast, the nuclear dataset (209,105 positions), which is ~10 times larger than the plastid dataset (~7.5 times larger if taking into account missing data, see Table 1), did not follow that expectation, with an average bootstrap support of 86%. The deep ochrophyte phylogeny inferred from the three compartments therefore showed not only incongruent relationships but also unexpected statistical supports.

### Comparison of the apparent phylogenetic signal across the three genomes when controlling for the number of sites and the number of OTUs

There was more apparent signal in the plastid than in the nuclear dataset for the deep branches connecting the major clades, PS’(cp|LG4X)>PS’(nu|LG4X), as shown by BS values (Fig. 1D). According to equation (2), this would indicate that λ_B_(cp)*EE(LG4X|cp,B) > λ_B_(nu)*EE(LG4X|nu,B), since n_cp_ ≪ n_nu_. However, differences in taxon sampling (64/63 species in the mitochondrial/plastid datasets versus 124 in the nuclear dataset) can affect extraction efficiency, making our comparison of the three datasets difficult to interpret. To cancel out the impact of taxon sampling on extraction efficiency, we reduced the sampling of each dataset to a common set of 23 species (22 for the mitochondrion). The phylogenies, inferred with the same LG4X model as above, (Supplementary Figure 4) were virtually identical to those inferred with more species (Supplementary Figures 1-3). Yet, we observed a slight decrease in the apparent phylogenetic signal when reducing the number of species: the average BS for the eight deep nodes went down from 73% to 64% for the mitochondrion, from 97% to 93% for the plastid and from 86% to 69% for the nucleus. This is in agreement with the widely recognized idea that the use of a large number of species improves phylogenetic accuracy, hence extraction efficiency (Zwickl and Hillis 2002). The higher apparent phylogenetic signal in the plastid versus nuclear compartment was thus still observed, in spite of controlling for taxon sampling (i.e., making EE(LG4X|D) closer for the three genomes).

To better characterize the apparent phylogenetic signal of the three compartments, we used the variable length bootstrap (VLB), or partial bootstrap, approach with the set of common species (Lecointre et al. 1994; Springer et al. 2001; Baurain et al. 2010). Usually, VLB analyses are used to define the number of sites needed to reach a predefined level of apparent phylogenetic signal (e.g., BS=95%), in order to compare the resolving power of different datasets (Springer et al. 2001; Baurain et al. 2010). Here, they allowed us to study the variation in apparent phylogenetic signal between the three compartments without being affected by the different sizes of the datasets.

Interestingly, VLBs revealed that the apparent phylogenetic signal of most nodes was highly similar for the nucleus, the plastid and, to a lesser extent, the mitochondrion (Fig. 2). For the monophyly of the major clades (Fig. 2A-F), VLB curves always reached 100% BS below 1000 positions. For the five higher-level groupings (BB, PD, PX, PXR, and BB+PD) that were easily recovered with the three genomes (Fig. 1A-C), the curves displayed similar increasing trends between compartments (Fig. 2G-K). The mitochondrial dataset required more sites to reach a given BS, which could be due to a reduced extraction efficiency related to the high rate of evolution in this compartment (Neiman and Taylor 2009). Yet, nucleus and plastid curves were virtually identical, sometimes the plastid increasing slightly faster (Fig. 2H) or slower (Fig. 2I) than the nucleus. In sharp contrast, support for the monophyly of E+SSC (Fig. 2L), as well as their subsequent grouping with Pinguiophyceae and PXR (Fig. 2M), rose much faster and higher in the plastid dataset than in the two other compartments. None of the bipartitions conflicting with E+SSC (Fig. 2N-P) showed the same rapid increase in the mitochondrion or the nucleus, showing that a strong apparent phylogenetic signal only exists in the plastid dataset for positioning these taxa.

**Figure 2.**
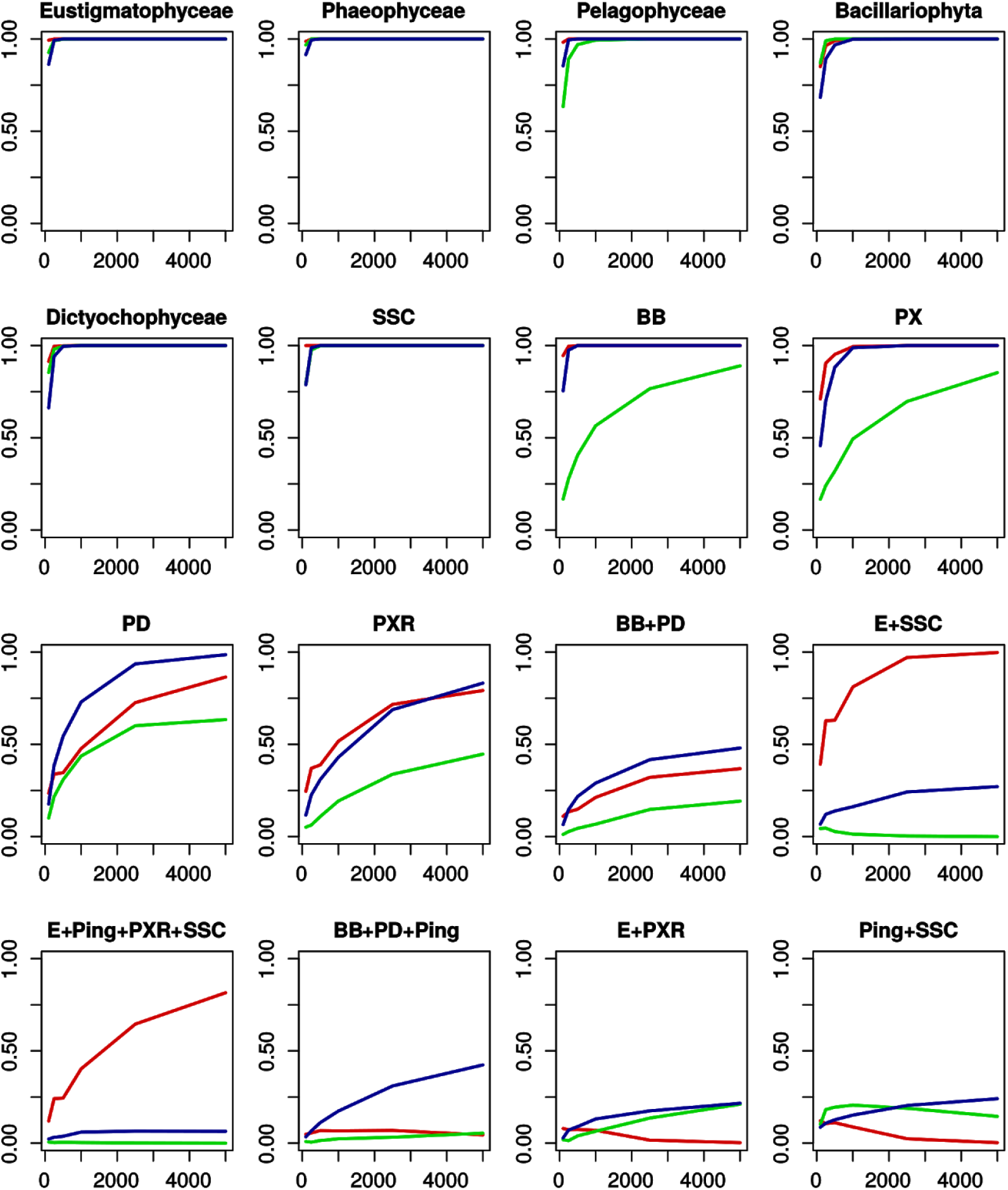
Variable length bootstrap results for a set of groupings in the three compartments under the LG4X model. X-axes represent the number of sites used to infer phylogeny, whereas Y-axes represent the bootstrap support observed for the grouping of interest. Line colors represent the compartments: nucleus (blue), plastid (red) and mitochondrion (green). (SSC) Synchromophyceae, Synurophyceae and Chrysophyceae, (BB) Bolidophyceae and Bacillariophyta, (PX) Phaeophyceae and Xanthophyceae, (PD) Pelagophyceae and Dictyochophyceae, (PXR) Phaeophyceae, Xanthophyceae and Raphidophyceae (E) Eustigmatophyceae, (Ping) Pinguiophyceae.

### Hypotheses to explain the conflict between plastid and nucleus

When controlling for the number of species and the number of sites, the apparent phylogenetic signal in plastid and nuclear compartments was almost identical for most nodes, meaning that λ_B_(cp)*EE(LG4X|cp,B) ~= λ_B_(nu)*EE(LG4X|nu,B). The higher average support observed over the eight deep nodes with the plastid dataset (Fig. 1D) was therefore due to a few nodes (e.g., E+SSC and E+Ping+PXR+SSC), for which λ_B_(cp)*EE(LG4X|cp,B) > λ_B_(nu)*EE(LG4X|nu,B). The comparison of branch lengths in Fig. 1 allowed us to formulate two hypotheses to account for this inequality. In the plastid tree (Fig. 1B), Eustigmatophyceae and SSC are connected by a long internal branch, which is ~3 times longer than the internal branch basal to Pinguiophyceae and SSC in the nucleus tree (Fig. 1C) and evolved much faster than the other clades. The first hypothesis is that E+SSC is correct and that this grouping benefits from a genuinely high value of λ_E+SSC_(cp). In contrast, the second hypothesis is that E+SSC is incorrect and that their grouping is the result of a long branch attraction (LBA) artifact, amounting to a very negative value of EE(LG4X|cp,B). Note that this is not the LBA artifact originally described by Felsenstein (1978) in the case of maximum parsimony, because probabilistic methods used here do take branch lengths into account. Nevertheless, fast evolving lineages not only evolve faster but also evolve differently from other lineages, being more subject to heterotachy (e.g., differences in the sets of sites free to vary (Lockhart 1996; Germot and Philippe 1999)) and/or heteropecilly (different substitution processes at play) (Roure and Philippe 2011), which violate the stationarity assumption made by almost all models. For the sake of simplicity, in what follows, we will present our results in terms of LBA, without expliciting anew that LBA in a probabilistic setting is due to model violations.

For illustrative purposes, let us make a simplifying assumption: either the plastid or the nucleus tree is fully representative of the true phylogeny (Fig. 3). In both cases, the durations between speciation events at the base of Ochrophyta are very short (on the left of Fig. 3). In the Plastid-Correct (PC) hypothesis (Fig. 3A), the long branch length of E+SSC is genuine in the plastid dataset (due to an increased evolutionary rate), hence a large λ_E+SSC_(cp). In addition, as both E and SSC evolved faster, LBA further favors a high extraction efficiency for this grouping (EE(LG4X|cp,E+SSC) > 1), which leads to the correct topology being inferred with very high bootstrap support. In contrast, in the nucleus tree, Pinguiophyceae and SSC evolved faster than E and PXR. Therefore, LBA between Pinguiophyceae and SSC creates a non-phylogenetic signal in favor of the erroneous groupings P+SSC and E+PXR (i.e., EE(LG4X|nu,E+SSC) < 0). In the Nucleus-Correct (NC) hypothesis (bottom of Fig. 3), E and SSC evolved much faster than the other ochrophytes in the plastid dataset, generating a strong LBA artifact (*e.g.*, correspondingly a very negative value for EE(LG4X|cp,P+SSC)) that exceeds the genuine P+SSC signal. As a result, λ_E+SSC_(cp)*EE(LG4X|cp,E+SSC) ≫ λ_P+SSC_(cp)*EE(LG4X|cp,P+SSC). The nucleus tree was easier to infer, because the fast evolving taxa (P and SSC on one hand and PD and BB on the other) are sister groups, so LBA reinforces the genuine phylogenetic signal. Because both hypotheses imply an erroneous branching due to a negative value of extraction efficiency (EE), distinguishing between them requires estimating whether the unavoidable model violations are sufficient to generate erroneous trees with the plastid or the nucleus datasets. Here, we applied two commonly used approaches against the LBA artifact (i.e., to reveal the effect of model violations): varying taxon sampling (to favor or disfavor LBA) and using different models of sequence evolution (more or less sensitive to the aforementioned model violations).

**Figure 3.**
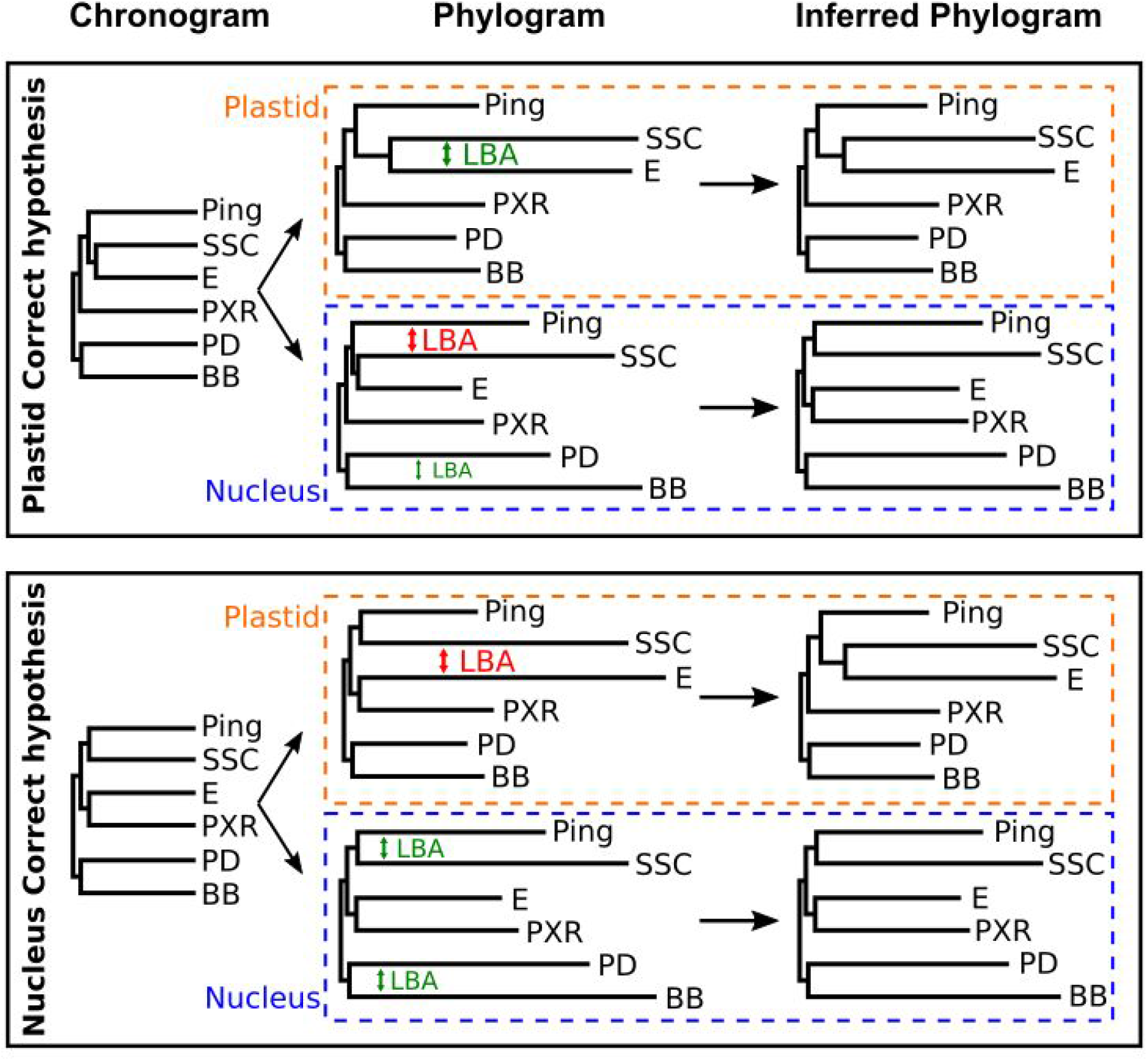
Two hypotheses to explain the incongruence between plastid and nucleus phylogenies. The first column shows the chronogram assumed to be correct in each hypothesis (top: Plastid-Correct, bottom: Nucleus-Correct), whereas the second column shows the corresponding phylograms (with branch lengths further accounting for the evolutionary rate of each lineage) and the third column the phylograms expected to be inferred when using a model unable to deal with LBA artifacts. Phylograms are shown for each compartment (top: plastid, bottom: nucleus). Depending on the true topology and the respective evolutionary rates of the lineages, LBA can either reinforce (green) or overwhelm (red) the phylogenetic (historical) signal, which results in incongruent apparent signals.

### Evidence for the presence of model violations

First, we evaluated the impact of major variations of the taxon sampling. The rationale was to reveal a possible inconsistency of the tree reconstruction method (i.e., model violations sufficiently important to make EE(m|D,B)<0) through the discovery of incongruence between phylogenies inferred from different subsets of species. We investigated two strategies: (1) use of only a distant outgroup (to increase LBA by creating a long unbroken branch) and (2) independant removal of highly supported ochrophyte lineages. We selected the six clades that were strongly supported by the three datasets: Eustigmatophyceae (E), Pinguiophyceae (Ping), SSC clade (represented by SC in the plastid), PXR clade, PD clade and BB clade. Since the phylogenetic signal is more accurately extracted with many taxa, we focus on the analyses with complete (albeit different, see above) taxon sampling. Analyses with the common set of 23 species returned comparable results, but with weaker BS values (Supplementary Table 1). All groupings of the six major clades observed through the 14 taxon sampling variations (2 compartments * 1+6 taxon samplings) are reported in Table 2.

**Table 2.** Bootstrap support of high-level ochrophyte clades with varying taxon sampling under the LG4X model

| Groupings | Plastid |  |  |  |  |  |  |  | Nucleus |  |  |  |  |  |  |  |
| --- | --- | --- | --- | --- | --- | --- | --- | --- | --- | --- | --- | --- | --- | --- | --- | --- |
|  | All | Out | E | SSC | Ping | PXR | PD | BB | All | Out | E | SSC | Ping | PXR | PD | BB |
| BB+E+PD+PXR |  |  | . |  |  | . | . | . |  | **78** | . |  |  | . | . | . |
| BB+E+PD+PXR+SSC |  |  | . |  |  | . | . | . |  | **100** | . |  |  | . | . | . |
| **BB+PD** | 100 | 100 | 98 | 90 | 100 | 99 | . | . | 68 | 100 | 97 |  | 100 |  | . | . |
| BB+PD+Ping |  |  |  |  | . |  | . | . |  |  |  | **95** | . | **88** | . | . |
| BB+PD+Ping+PXR |  | **95** |  |  | . | . | . | . |  |  |  |  | . | . | . | . |
| BB+PD+PXR |  | **98** |  |  | **56** | . | . | . |  |  |  |  |  | . | . | . |
| BB+Ping |  |  |  |  | . |  |  | . |  |  |  | **95** | . | **88** | **83** | . |
| E+Ping |  |  | . | *67* | . |  |  |  |  |  | . |  | . |  |  |  |
| E+Ping+PXR |  |  | . | *71* | . | . |  |  |  |  | . |  | . | . |  |  |
| E+Ping+PXR+SSC | 82 |  | . | . | . | . | *77* | *53* | 67 |  | . | . | . | . |  | 100 |
| E+Ping+SSC | 95 |  | . | . | . | 91 | 80 | 66 |  |  | . | . | . |  |  |  |
| **E+PXR** |  |  | . |  |  | . |  |  | 81 | 82 | . | 96 |  | . | 91 | 100 |
| E+PXR+SSC |  |  | . | . |  | . |  |  |  |  | . | . | *83* | . | **83** |  |
| **E+SSC** | 100 | 100 | . | . | 100 | 100 | 100 | 100 |  |  | . | . | **56** | **91** |  |  |
| Ping+PXR+SSC |  |  | *87* | . | . | . |  |  |  |  | 96 | . | . | . |  |  |
| **Ping+SSC** |  |  | 95 | . | . |  |  |  | 68 |  | 97 | . | . |  |  | 100 |

For the plastid, only two taxon sampling variations produced an incongruent topology, the use of a distant outgroup and the removal of Pinguiophyceae (3 incompatibilities, BS shown in boldface in Table 2). Both resulted in the same artifactual topological move: the attraction of the fast-evolving E+SSC group by the outgroup. Attractions were explained by the presence of a longer unbroken branch, either the distant outgroup or the branch of E+SSC in the absence of their slowly-evolving sister-group Pinguiophyceae. Taxon sampling variations of the plastid dataset revealed that model violations of LG4X sometimes produced LBA artifacts. This indicated a limited extraction efficiency with this combination of model and dataset, suggesting that the hypothesis NC may be correct. Yet, it is important to notice that the grouping E+SSC was always observed.

Variations of the taxon sampling in the nucleus dataset showed more incompatibilities with the tree inferred from all species (10, in boldface in Table 2, corresponding to 6 alternative groupings) than in the plastid (only 3). The incompatibilities were more complicated to understand, as the six clades evolved at a more homogeneous rate in the former than in the latter (Fig. 1B-C). Pinguiophyceae appeared to be the most unstable clade: they emerged as the sister-group to the remaining ochrophytes (BS=100%) when using the distant outgroup, as the sister-group to BB (BS=95%) when removing SSC, and as the sister-group to SSC (BS=100) when removing BB. Pinguiophyceae were only represented by two closely related species (*Phaeomonas* and *Pinguiococcus*), leaving a long unbroken branch at their base (Fig. 1C). As BB and SSC are the fastest evolving clade in the nuclear dataset, the placement of Pinguiophyceae can be explained by a LBA with the longest branch available in each of the three cases, *i.e.*, the outgroup, BB, and SSC, respectively. The limited support for Ping+SSC (BS=68%) when using the complete dataset could then result from an average among these contradictory attractions, all the more so that the competing bipartition (32%) is BB+Ping. The removal of PXR, a relatively slowly evolving clade, had the most dramatic effect, all the deep relationships becoming incongruent. It may have allowed the grouping of E+SSC (BS=91%), hence reducing the attraction between SSC and Pinguiophyceae, the latter being then attracted by BB (BS=88%). Interestingly, E+SSC was also recovered through the removal of Pinguiophyceae (BS=56%). Altogether, these results suggest the presence of a high amount of non-phylogenetic signal under the LG4X model and/or a limited genuine phylogenetic signal in the nuclear dataset (i.e., a limited extraction efficiency), thereby supporting hypothesis PC.

While taxon sampling variations of the nuclear dataset argued in favor of hypothesis PC, as E+PXR and Ping+SSC groupings failed to be robustly recovered, the plastid dataset also showed incongruence that may instead argue in favor of hypothesis NC. The higher number of incompatibilities observed with the nucleus than the plastid (10 versus 3) indicates an extraction efficiency and/or an amount of genuine phylogenetic signal that are lower with the former (*i.e.*, EE(LG4X|nu,B) < EE(LG4X|cp,B) and/or λ_B_(nu) < λ_B_(cp)) and yields a less reliable tree. For instance, whereas plastid taxon samplings consistently recovered two high-level clades (E+SSC and BB+PD), the nuclear dataset failed to recover any such clade consistently. However, the main result of taxon sampling variations was the evidence for a major impact of model violations with LG4X, especially for the nuclear compartment. These observations are in agreement with the sensitivity to LBA of models that do not fully incorporate the heterogeneity of the substitution process across sites (Lartillot et al. 2007; Philippe et al. 2011; Simion et al. 2017). They further suggest that neither the PC nor the NC hypothesis is correct and that we need to use a better model to get insights into the ochrophyte radiation.

### Impact of using a better fitting model of sequence evolution

We tested two site-heterogeneous models that have been shown to be less sensitive to LBA (Lartillot et al. 2007): the CAT model implemented in the Bayesian framework (Lartillot and Philippe 2004) and the C20+LG model, an empirical version of the CAT model implemented in the maximum likelihood format (Le et al.. 2008). We first compared LG4X to C20+LG using ModelFinder (Kalyaanamoorthy et al. 2017) from IQTREE (Nguyen et al. 2015), which showed C20+LG to be better than LG4X with both the plastid and the nuclear datasets (Supplementary Table 2). Second, cross-validation demonstrated CAT to have a better fit than C20+LG for both datasets (plastid: likelihood difference between CAT and C20+LG of 370 +/− 152; nucleus: 488 +/− 141). Consequently, the combination of these two tests showed CAT to have a better fit than LG4X and C20+LG for our datasets. Even if CAT is computationally very demanding (*e.g.*, Philippe et al. 2019), we were able to compute CAT bootstrap support for the complete plastid dataset given its moderate size (63 × 21,692). In contrast, this computationally expensive model could not be used on the much larger nuclear dataset (124 × 209,105). To make the analysis tractable, we resorted to a gene jackknife approach instead (Delsuc et al. 2008). We chose to generate datasets of ~50,000 positions and to run 50 replicates. In the case of LG4X, we verified that the jackknife supports (JS) were comparable with BS, despite being based on ~4 times less positions: as expected, JS values were lower than BS values (Supplementary Table 3). Yet, and more importantly, the same groupings were recovered in all but one case (the position of SSC when using a distant outgroup). Therefore, JS is a reasonable proxy for BS to evaluate the effect of taxon sampling on the nucleus-based phylogeny inferred with the better fitting CAT model.

The plastid tree inferred using the CAT model (Fig. 4A) had the same topology as the LG4X tree, but with lower statistical support, especially for E+SSC (BS=84% versus 100%) and the position of its sister-group Pinguiophyceae (BS=58% versus 95%). Lower support for E+SSC can be explained by the fact that the CAT model is more suspicious when it has to group two long branches (Eustigmatophyceae and SSC) together. In other words, it assumes more shared amino acids to be due to convergence than LG4X, the very reason for its reduced sensitivity to LBA (Lartillot et al. 2007). In contrast, the topology inferred from the nucleus supermatrix using CAT (Fig. 4B) was different from that inferred with LG4X (Fig. 1C): only the monophyly of BB+PD (JS=76%) was common among the high-level relationships observed between the six major clades. In the CAT tree, SSC was sister of Eustigmatophyceae (E+SSC; JS=54%) instead of Pinguiophyceae, while the latter group was sister of BB+PD (JS=56%). Finally, PXR was weakly grouped with BB+PD+Ping (JS=32%). Overall, the use of a better fitting model, which likely improves extraction efficiency, allows the common recovery of the relationship between Eustigmatophyceae and SSC (E+SSC) by both the plastid and the nucleus datasets, a relationship key to distinguish between the two hypotheses explaining the conflicts observed between the two compartments when using the LG4X model (Fig. 3).

**Figure 4.**
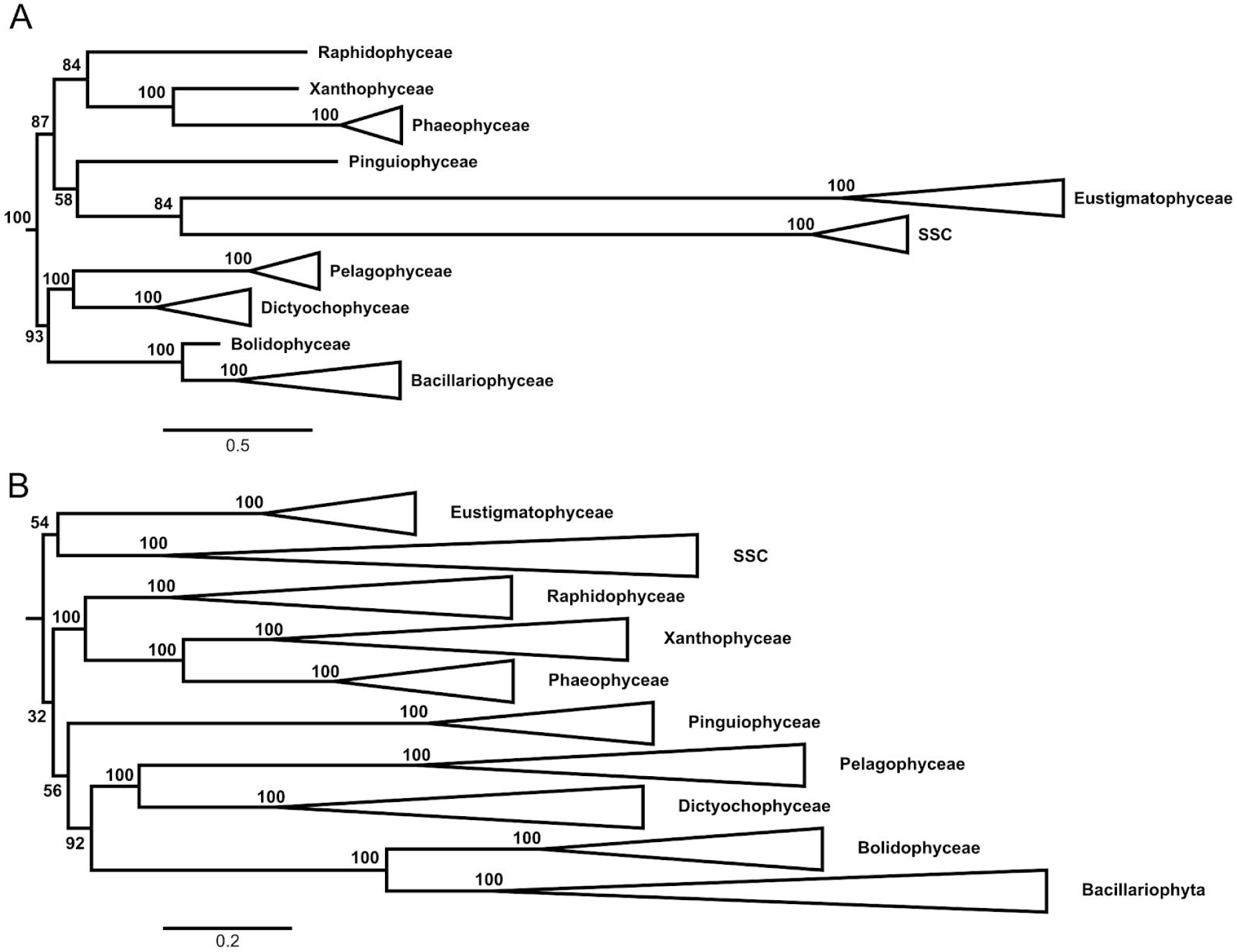
Phylogenetic trees inferred using PhyloBayes-MPI under the CAT+Γ4 model. The ten major clades of Ochrophyta have been collapsed when more than two species were present. Statistical support values are displayed next to their relative nodes. (A) Plastid dataset (63 species and 21,692 positions). Statistical support based on 100 non-parametric bootstrap replicates. (B) Nuclear dataset (124 species and 209,105 positions). Statistical support based on 50 gene jackknife replicates of about 50,000 positions. Interestingly, the three most frequent alternative groupings for the nucleus, Ping+SSC (40%), E+PXR (20%) and E+Ping+PXR+SSC (20%), were all recovered with LG4X (Fig. 1C).

To confirm that using a better fitting model increases extraction efficiency and therefore reduces incongruence, we performed the same variations of taxon sampling as above, but using the CAT model. The results (Table 3) clearly revealed less incompatibilities within each compartment (1 versus 3 for the plastid and 4 versus 10 for the nucleus) and better congruence between the two compartments. In particular, CAT recovered BB+PD and E+SSC in all analyses of the two compartments. The position of Pinguiophyceae and PXR remained unstable, displaying various sister relationships to one of the two previous clades with limited support. However, the nucleus supported the relationship between Pinguiophyceae and BB+PD in all taxon sampling experiments, except when Eustigmatophyceae were removed, in which case Pinguiophyceae were sisters to fast-evolving SSC, hence possibly a LBA artifact. In contrast, the plastid dataset did never recover BB+PD+Ping. Despite the use of fewer sites (50,000) than LG4X (i.e., increased stochastic error), CAT thus turned out to be more robust to taxon sampling variations, thereby demonstrating its success in increasing extraction efficiency (i.e., reducing the amount of non-phylogenetic signal).

**Table 3.**
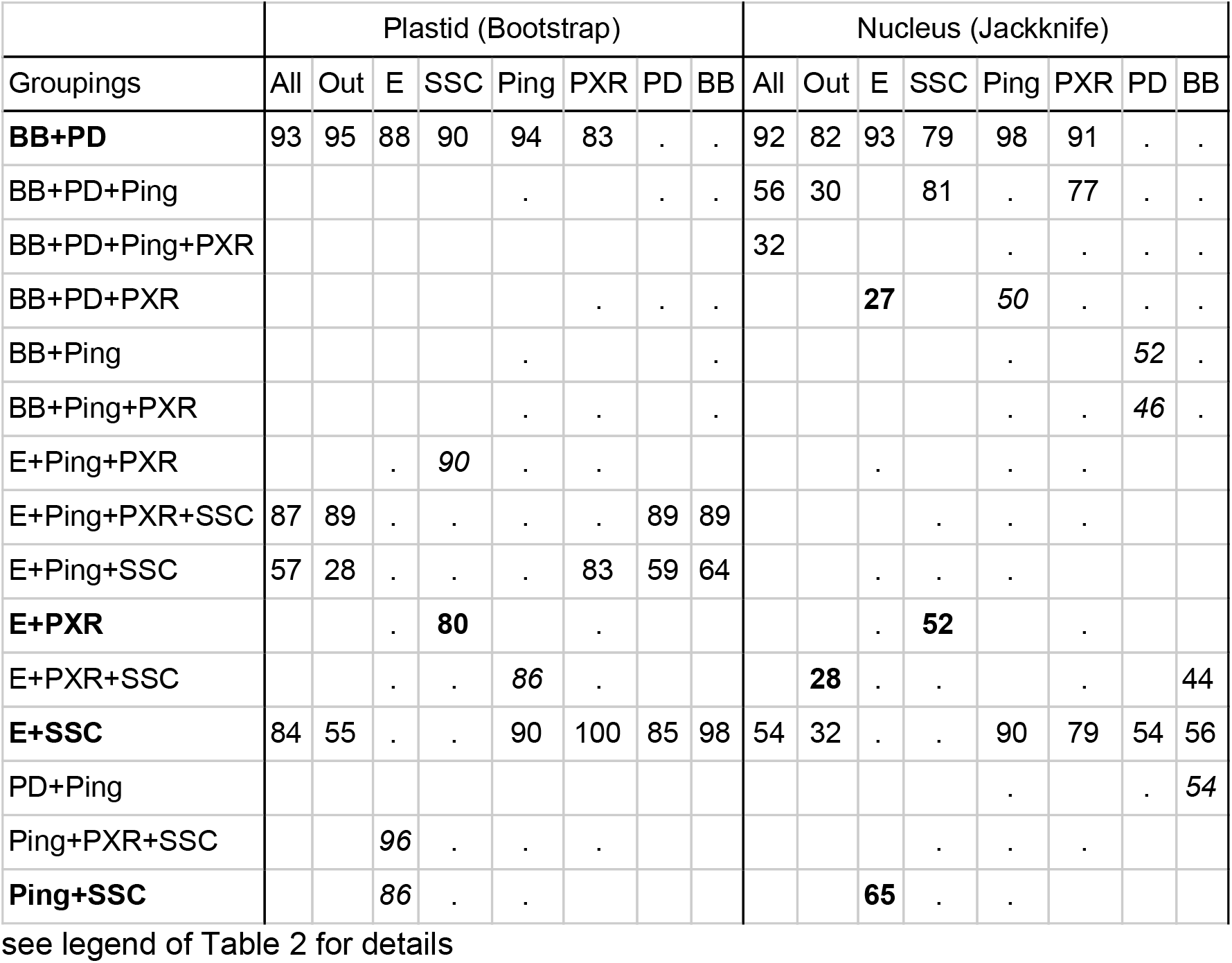
Support of high-level ochrophyte clades with varying taxon sampling under the CAT+Γ4 model

| Groupings | Plastid (Bootstrap) |  |  |  |  |  |  |  | Nucleus (Jackknife) |  |  |  |  |  |  |  |
| --- | --- | --- | --- | --- | --- | --- | --- | --- | --- | --- | --- | --- | --- | --- | --- | --- |
|  | All | Out | E | SSC | Ping | PXR | PD | BB | All | Out | E | SSC | Ping | PXR | PD | BB |
| **BB+PD** | 93 | 95 | 88 | 90 | 94 | 83 | . | . | 92 | 82 | 93 | 79 | 98 | 91 | . | . |
| BB+PD+Ping |  |  |  |  | . |  | . | . | 56 | 30 |  | 81 | . | 77 | . | . |
| BB+PD+Ping+PXR |  |  |  |  |  |  |  |  | 32 |  |  |  | . | . | . | . |
| BB+PD+PXR |  |  |  |  |  | . | . | . |  |  | **27** |  | 50 | . | . | . |
| BB+Ping |  |  |  |  | . |  |  | . |  |  |  |  | . |  | 52 | . |
| BB+Ping+PXR |  |  |  |  | . | . |  | . |  |  |  |  | . | . | 46 | . |
| E+Ping+PXR |  |  | . | 90 | . | . |  |  |  |  | . |  | . | . |  |  |
| E+Ping+PXR+SSC | 87 | 89 | . | . | . | . | 89 | 89 |  |  |  | . | . | . |  |  |
| E+Ping+SSC | 57 | 28 | . | . | . | 83 | 59 | 64 |  |  | . | . | . |  |  |  |
| **E+PXR** |  |  | . | **80** |  | . |  |  |  |  | . | **52** |  | . |  |  |
| E+PXR+SSC |  |  | . | . | 86 | . |  |  |  | **28** | . | . |  | . |  | 44 |
| **E+SSC** | 84 | 55 | . | . | 90 | 100 | 85 | 98 | 54 | 32 | . | . | 90 | 79 | 54 | 56 |
| PD+Ping |  |  |  |  |  |  |  |  |  |  |  |  | . |  | . | 54 |
| Ping+PXR+SSC |  |  | 96 | . | . | . |  |  |  |  |  | . | . | . |  |  |
| **Ping+SSC** |  |  | 86 | . | . |  |  |  |  |  | **65** | . | . |  |  |  |

see legend of Table 2 for details

We then estimated the performance of the computationally more efficient, but less fitting, site-heterogeneous model C20+LG. As expected from its intermediate fit between LG4X and CAT, the impact of taxon sampling (Supplementary Table 4) was intermediate: 4 incongruences for the plastid and 3 for the nucleus with C20+LG (to compare with 3 and 10 for LG4X and 1 and 4 for CAT). Importantly, only one grouping, BB+PD, was consistently recovered in all experiments. While E+SSC was always recovered by the plastid dataset, the nucleus dataset found it in only one case (after removal of PXR) and generally grouped Eustigmatophyceae with PXR and SSC with Pinguiophyceae. Albeit less sensitive to taxon sampling, C20+LG did not improve the congruence between the plastid and the nucleus, suggesting that its model violations remained serious. As a result, despite C20+LG can handle the complete dataset, the site-heterogeneous CAT model appeared as the most suitable to accurately address the difficult question of the Ochrophyta radiation.

Improving extraction efficiency favored one facet of hypothesis PC (Fig. 3), *i.e.*, the grouping of Eustigmatophyceae and SSC is correct and more highly supported in the plastid than in the nucleus compartment because of an acceleration of the substitution rate (and thus of λ_E+SSC_(cp)) in the branch at the base of E+SSC in plastid loci. First, E+SSC was always recovered by the two compartments for sixteen (2*8) different taxon sampling variations. Second, this relationship is also supported by a common split of the plastid encoded gene clpC, which is involved in the protein degradation pathway mediated by the ClpP protease (Ševčíková et al. 2015). Third, we observed five common losses of plastid genes in these two clades (ATP synthase CF1 delta subunit, hypothetical protein Ycf39/Isoflavone reductase, PSI reaction center subunit XII, hypothetical protein Ycf35 and cytochrome b6-f complex subunit 6/petL; data not shown), although we cannot exclude convergence since the plastid genome is reduced in both cases. Overall, our experiments proved that improving extraction efficiency through selection of a better fitting model reduced incongruence and increased robustness to taxon sampling variations. Albeit only two (BB+PD and E+SSC) high-level relationships out of four were consistently recovered in the case of Ochrophyta, the use of the CAT model demonstrated the key importance of the extraction efficiency in resolving ancient radiations.

### Using branch length heterogeneity to tackle radiations

Hypothesis PC (Fig. 3) postulates an acceleration of the evolutionary rate in the (genuine) internal branch connecting Eustigmatophyceae and SSC in the plastid compartment (i.e., high value of λ_E+SSC_(cp)). However, owing to the important model violations that favor LBA with LG4X (Table 2), the grouping of the fast evolving Eustigmatophyceae and SSC obtained with this model (and the corresponding internal branch length) is likely inflated (i.e., EE(LG4X|cp,E+SSC)>1). It is thus difficult to assess the relative value of λ_E+SSC_(cp) versus λ_E+SSC_(nu). Even with the better fitting CAT model, model violations can inflate the apparent phylogenetic signal for E+SSC. If we neglect the confounding effect of model violations in estimating the true value of λ_E+SSC_, this underlines the interest of finding markers with relatively long internal branches (i.e., possible high value of λ) to resolve radiations.

Obviously, looking for such genes is difficult because it requires being able to accurately infer the value of λ. However, testing the potential of such an approach is possible by assuming the knowledge of the correct phylogeny. More precisely, we can estimate branch lengths for each gene, constrained to a candidate topology, and select those displaying the longest (or shortest) length for the branch of interest. Finally, we can infer a phylogeny using a concatenation of the resulting set of markers and compare it to the phylogeny obtained without such a selection to study the effect of filtering the dataset by the signal of interest.

We applied this protocol, using the LG4X model, to the nucleus dataset by selecting the 200 genes with the longest internal branch at the base of E+SSC, yielding a supermatrix of 47,386 positions (LONG_nu_) and, as a negative control, the 200 genes with the shortest internal branch, yielding a supermatrix of 39,867 positions (SHORT_nu_). Not surprisingly, the phylogeny inferred from SHORT_nu_ with the LG4X model (Supplementary Figure 5A) did not recover E+SSC, but strongly grouped Pinguiophyceae and SSC (BS=96%) and Eustigmatophyceae and PXR (BS=100%), in agreement with the topology observed with the full dataset (Fig. 1C). In the absence of a strong genuine phylogenetic signal (for E+SSC), the misleading non-phylogenetic signal dominated, and the apparent phylogenetic signal for two erroneous groupings (P+SSC and E+PXR) increased (BS rose from 68/81 to 96/100, respectively). Note that a zero branch length might also be due to the fact that a locus has a different history (*e.g.*, due to hybridization or ILS), amounting to the branch being non-existent. When the time separating two nodes is very short, the probability to observe at least one substitution in the corresponding branch is proportional to the size of the genes. We therefore expect the genes having a very short branch length to be shorter in terms of positions than the ones with a long branch. This prediction is fulfilled (199 versus 237 positions on average), suggesting that rate variation and short gene length rather than different history are the main causes of the observed short branch lengths.

In contrast, the phylogeny inferred from LONG_nu_ with LG4X (Supplementary Figure 5B) strongly supported E+SSC (BS=100%) with the same complete taxon sampling. This suggests that the genuine phylogenetic signal was now stronger than the non-phylogenetic signal created by the serious violations affecting this model, thereby leading to a strong apparent signal in favor of E+SSC. As the full dataset did not support E+SSC, under the assumption that extraction efficiency of LG4X is the same for all genes, the non-phylogenetic signal produced over 209,105 positions is probably stronger than the corresponding signal in the 47,386 positions of the LONG_nu_ set of genes. This protocol cannot be used to resolve radiations, because it assumes the species phylogeny to be known, but it can be used to reveal the contradictory attractions present in a large dataset (here SSC attracted either by Pinguiophyceae or Eustigmatophyceae), these attractions stemming either from model violations or from the genuine (historical) signal. More importantly, it validates the idea of looking for innovative methods to detect genes with a high signal for internal nodes of a species phylogeny, disregarding the global topologies of the gene trees. Such approaches could be another avenue to alleviate the impact of model violations when trying to resolve radiations, without designing ever-more complex evolutionary models.

### Towards resolving the Ochrophyta phylogeny

Our analysis showed that the resolution of the deep ochrophyte relationships was extremely difficult, because of short internal branches and serious model violations. Interestingly, the small plastid dataset appeared to contain a relatively large amount of phylogenetic signal, in particular because of its high value of λ_E+SSC_(cp). Since the CAT model did not show evidence of a strongly negative value of extraction efficiency for the nucleus or the plastid, it should be interesting to combine the high number of positions of the nucleus and the high λ of the plastid to increase the apparent phylogenetic signal of the ochrophyte radiation. Indeed, by combining the plastid and nuclear datasets, we should lengthen at least one of the difficult branches, i.e., λ_E+SSC_(nu+cp) > λ_E+SSC_(nu), thus making the problem easier to resolve for any method of phylogenetic inference. However, there are potential drawbacks to this approach, such as the fact that combining those datasets would reduce the taxon sampling down to 23 common species, along with the potential introduction of additional model violations (in particular branch length heterogeneity across compartments) (Kolaczkowski and Thornton 2004). Whereas these drawbacks might both decrease extraction efficiency, reducing the number of species allowed us to use more sites (80,000) with the best fitting model (CAT).

Such a combined phylogeny inferred with the nu+cp supermatrix using the CAT model (Fig. 5) showed a much higher support for deep ochrophyte relationships: BB+PD (JS=100%), BB+PD+Ping (JS=100%), E+SSC (JS=99%) and E+PXR+SSC (JS=90%). Interestingly, the nu+cp phylogeny (Fig. 5) is different from both the plastid (Fig. 4A) and the nucleus (Fig. 4B) trees. However, a higher statistical support is not necessarily incontrovertible evidence for a given grouping, as the inference method might be inconsistent. Therefore, we applied the same taxon sampling variations as above (*i.e.*, the use of a distant outgroup and the removal of each major ochrophyte clade) to the nu+cp supermatrix. Interestingly, all seven variations returned trees fully compatible with the phylogeny of Fig. 5 (Table 4). In contrast, the use of a less fitting model (LG4X) on the same supermatrix yielded a lower support and displayed sensitivity to taxon sampling (Supplementary Table 5), thereby confirming the key role of the model of sequence evolution in the accurate resolution of short internal branches. Even under difficult phylogenetic inference conditions (limited extraction efficiency due to a small number of taxa and residual violations affecting the CAT model), the robustness to taxon sampling variations argued for the nu+cp phylogeny (Fig. 5) to be a credible working hypothesis for the deep ochrophyte relationships.

**Figure 5.**
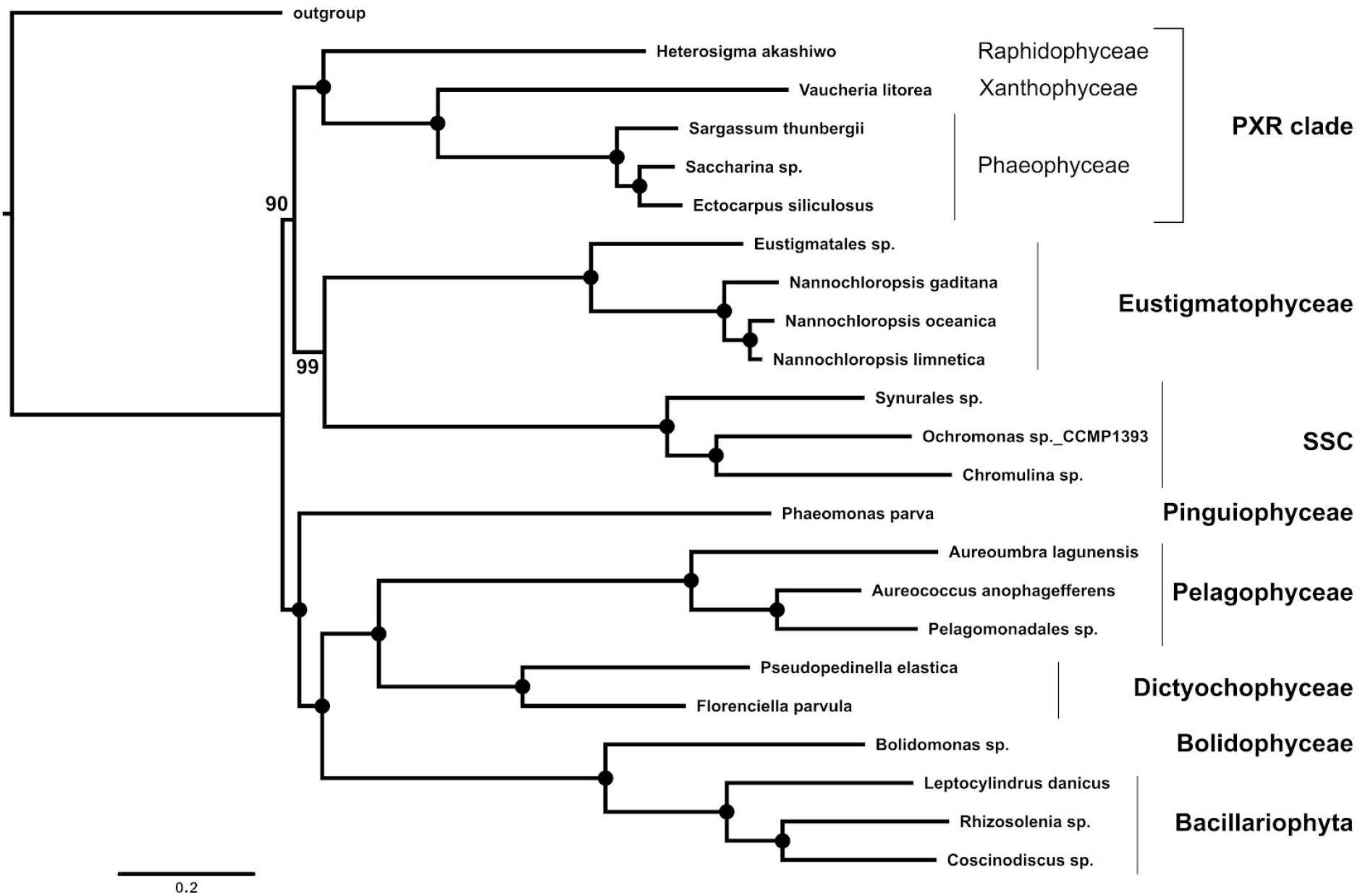
Consensus phylogenetic tree of the fusion (nu+cp) dataset, inferred from 100 jackknife replicates (~80,000 positions) under the CAT+Γ4 model using PhyloBayes-MPI. Statistical support corresponds to jackknife support (JS), with black circles meaning 100% JS. Species named *sp.* correspond to chimeras between the corresponding species of the plastid and nuclear dataset presented in Supplementary Table 5.

**Table 4.**
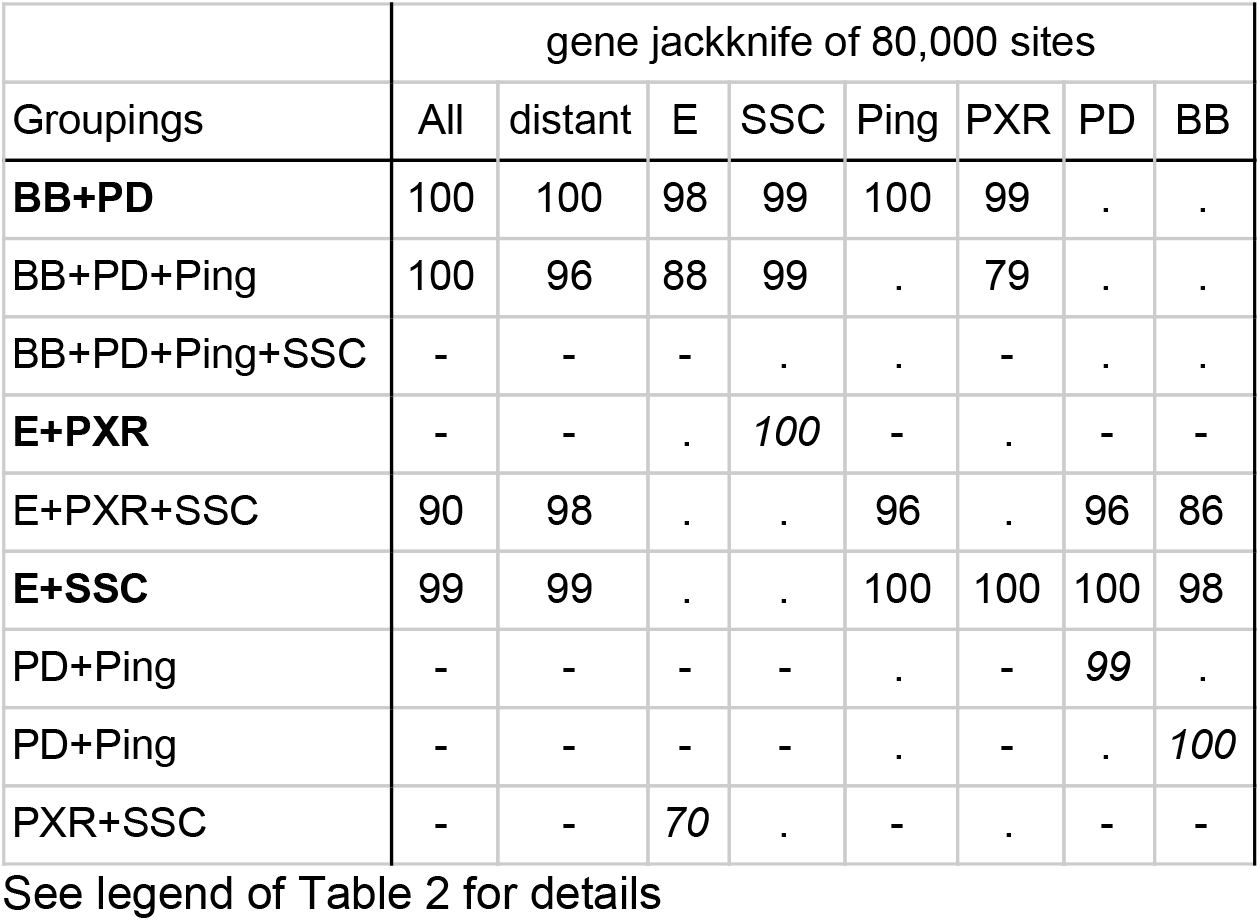
Jackknife support of high-level ochrophyte clades of the fusion (nu+cp) dataset with varying taxon sampling under the CAT+Γ4 model

|  | gene jackknife of 80,000 sites |  |  |  |  |  |  |  |
| --- | --- | --- | --- | --- | --- | --- | --- | --- |
| Groupings | All | distant | E | SSC | Ping | PXR | PD | BB |
| **BB+PD** | 100 | 100 | 98 | 99 | 100 | 99 | . | . |
| BB+PD+Ping | 100 | 96 | 88 | 99 | . | 79 | . | . |
| BB+PD+Ping+SSC | - | - | - | . | . | - | . | . |
| **E+PXR** | - | - | . | 100 | - | . | - | - |
| E+PXR+SSC | 90 | 98 | . | . | 96 | . | 96 | 86 |
| **E+SSC** | 99 | 99 | . | . | 100 | 100 | 100 | 98 |
| PD+Ping | - | - | - | - | . | - | 99 | . |
| PD+Ping | - | - | - | - | . | - | . | 100 |
| PXR+SSC | - | - | 70 | . | - | . | - | - |

See legend of Table 2 for details

## Conclusion

A common belief is that increasing the number of positions (n) has the potential to resolve evolutionary radiations. Our work confirms that this is a necessary condition (Fig. 2 and Supplementary Table 6), but that the two other components of the apparent phylogenetic signal formula proposed here, i.e., branch length (λ) and extraction efficiency (EE), cannot be neglected. In particular, extraction efficiency is a major limiting factor, because our models are necessarily over-simplified with respect to the complexity of biological evolution. The accumulation of data, not of positions but of species, is certainly useful, as the use of more taxa generally helps in the extraction of the phylogenetic signal. Yet, this approach has some serious limitations: (1) some branches are unavoidably unbroken because of extinction, (2) some (rogue) species decrease extraction efficiency, and (3) the resulting increase in computational time limits us to the use of the simplest models. Studies are thus needed to evaluate what are the best compromises between the number of species and the complexity of models to optimize extraction efficiency.

The reduction of model violations achieved when dropping LG4X in favor of the CAT model allowed us to reduce the incongruence revealed by taxon sampling variations and improve the resolution of the ochrophyte radiation, especially for the nucleus dataset. However, the CAT model is still far from perfect. For instance, it does not take into account the genetic code to weigh amino acid substitutions (see Rodrigue et al. 2010), and it assumes that the evolutionary process is the same all over the phylogeny (e.g., ignoring compositional biases, heterotachy or heteropecilly). These simplifications are bound to result in model violations that could lead to an incorrect phylogeny. The improvement of models of sequence evolution, both in terms of fit and of computational efficiency, should thus be a priority to resolve ancient radiations. For recent radiations, the impact of these model violations is expected to be more limited (fewer multiple substitutions at the same position), and it is key to address another kind of model violation (not studied in our work), the presence of inter-species gene flux (hybridization) and ILS, using coalescent methods such as *BEAST (Heled and Drummond 2010). However, when a non-negligible fraction of gene trees is different from the species tree, the interest of resolving the radiation is limited because hemiplasy is so frequent that the species tree is no longer useful to study the evolution of characters and organisms (Hahn and Nakhleh 2016).

In addition to the number of positions and the extraction efficiency, the strength of the apparent phylogenetic signal is also dependent on branch length. The length of a given branch is variable across loci, *e.g.*, being longer at a locus that underwent reduced purifying selection or directional selection. In the case of the plastid dataset, we were lucky to have had a large number of loci that underwent a substitution rate acceleration in the E+SSC basal branch. This acceleration likely explains the observation that the small plastid dataset (21,692 positions) is able to strongly recover the monophyly of this clade otherwise very difficult to resolve, whereas the large nuclear dataset (209,105 positions) cannot. The difference was more pronounced with the LG4X model than with the CAT model, probably because the long branches of Eustigmatophyceae and SSC were further artifactually attracted (i.e., EE(LG4X|cp,E+SSC)>1). This “lucky” rate acceleration suggests a new approach to resolve ancient radiations: searching for loci having accelerated in the short internal branches of interest, so as to facilitate extraction of a signal that is scarce for other, more regular, loci.

Finally, combining the nuclear and plastid datasets, along with the use of the CAT model, helped us to simultaneously increase n, λ and EE, leading to a well-supported tree, robust to taxon sampling variations. Given the difficulties to resolve the ochrophyte radiation, this phylogeny needs to be confirmed with a richer taxon sampling and/or with a better model. It nevertheless constitutes a working hypothesis to understand in which order the remarkably diverse phenotypes of Ochrophyta emerged, from the picoplanktonic *Nannochloropsis* to the silica frustule-bearing diatoms and to giant marine kelps.

## Materials and Methods

### Cultures, organelle genome sequencing and assembling

Cultures of *Chromulina chionophila* (CCAP 909/9), *Pseudopedinella elastica* (SAG B43.88), *Synura petersenii* (CCAC 0052), *Phaeomonas parva* (CCMP 2877), and *Florenciella parvula* (RCC 446) were obtained from their respective algal culture collections (CCAP: https://www.ccap.ac.uk/, SAG: http://www.uni-goettingen.de/en/culture+;collection+of+algae+%28sag%29/184982.html, CCAC: https://www.uni-due.de/biology/ccac/, CCMP: https://ncma.bigelow.org/, RCC: http://roscoff-culture-collection.org/). Algae were grown in the culture media recommended by the collections in aerated 1 L Erlenmeyer flasks at 15 °C and 20 μmol photons/m2/s in a 14:10 hr L/D cycle. They were harvested by centrifugation and, after grinding in liquid nitrogen, total DNA was extracted using either the NucleoSpin^®^ Plant II Midi Kit (Macherey-Nagel, Düren, Germany) or a modified CTAB protocol (Rogers and Bendich 1985; see Supplementary Information).

### Sequencing and assembly of organelle genomes

DNA samples were converted to Illumina sequencing libraries according to the manufacturer’s protocols and sequenced in paired end mode (150 bases sequencing length). The resulting reads were assembled using ABySS (Simpson et al. 2009). Organellar contigs were extracted using gene sequences from the respective *Ectocarpus* genomes as queries in BLAST searches. Gaps were closed with GapFiller (Nadalin et al. 2012) and annotation was carried out with the sequin tool from NCBI.

### Creation of phylogenomic datasets

For each compartment, we assembled the datasets following a semi-automatic protocol similar to the one described in our previous phylogenomic studies (Simion et al. 2017, Irisarri et al. 2017). In summary (see Figure 1 of Simion et al. for an overview and https://github.com/psimion/SuppData_Metazoa_2017/blob/master/utilities_src.tgz for software availability), we used protein annotations obtained from genomic data to define orthologous groups with OrthoFinder version 1.4 (Emms and Kelly 2015). Sequence similarity matrices were computed with BLAST for mitochondrial and plastid datasets and with USEARCH (Edgar 2010) for the nuclear dataset (e-value threshold = 1e-5) before being divided with the MCL algorithm using the default inflation value (1.5). We filtered the resulting orthogroups for minimal taxonomic representation before validating their orthology relationships. Then we improved their taxon sampling by adding species from transcriptomic and genomic data using Forty-Two (https://metacpan.org/release/Bio-MUST-Apps-FortyTwo). Detailed description for each compartment, as well as on the computational treatments undertaken to remove paralogous and xenologous sequences from the multiple sequence alignments, can be found in the Supplementary Information. In particular, we run the BLC method (Simion et al. 2017) to detect and remove outlier genes. Finally, our analyses focused on the three datasets summarised in Table 1 and available at **https://doi.org/10.6084/m9.figshare.7680395.v2**.

### Phylogenetic inferences

All supermatrices used in our analyses were concatenated using SCaFoS (Roure et al. 2007). We inferred phylogenetic trees using RAxML version 8.2 (Stamatakis 2014) with the LG4X mixture model (Le et al. 2012) using 100 fast bootstrap replicates. Inferences under the C20+LG model were carried out using IQ-TREE 1.6.8 (Nguyen et al. 2015). Inferences under the CAT+Γ4 mixture model (Lartillot and Philippe 2004) were carried out using PhyloBayes-MPI version 1.8 (Lartillot et al. 2013), either on bootstrap replicates for mitochondrial and plastid datasets or on gene jackknife replicates for the nucleus dataset. Preliminary analyses demonstrated that convergence was not reachable for a dataset of 124 species and 209,105 amino-acid positions with the current implementation of PhyloBayes-MPI. Following Delsuc et al. (2008), we thus used a gene jackknife approach and generated replicates of ~50,000 or ~80,000 positions with a custom script. Convergence assessment and consensus tree construction were performed as in Simion et al. (2017).

### VLB analyses

We reduced each dataset to an ingroup taxon sampling of 22 comparable species (21 for the mitochondrion as one out of four Pelagophyceae species was missing), *i.e.*, identical or closely related (Supplementary Table 7). For the outgroup, we used *Guillardia theta* for the plastid and *Phytophthora sojae* for the mitochondrion and *Phytophthora parasitica* for the nucleus. We used distinct species to have a similar branch length leading to the outgroup in each compartment, whereas using the same species (*e.g.*, *G. theta*) would have generated a much longer branch in the mitochondrion/nucleus than in the plastid. Out of the three resulting supermatrices, we drew 1000 variable length bootstrap (VLB) replicates of different sizes (100, 250, 500, 1000, 1500, 2000, 2500 sites) and 100 replicates of 5000 sites using seqboot from the PHYLIP package (Felsenstein 1989). The best tree was obtained for each VLB replicate with RAxML under the LG4X mixture model. Finally, we retrieved the bootstrap proportion of each bipartition for each matrix length with the program consense from the PHYLIP package, and further analyzed them using a custom R script.

### Model comparison

AIC, AICc and BIC between LG4X and GTR+Γ4 models were computed using ModelFinder (Kalyaanamoorthy et al. 2017) from IQTREE version 1.6.8 (Nguyen et al. 2015), with the constrained topology previously obtained under the LG4X model with RAxML. Cross-validations between GTR+Γ4 and CAT+Γ4 were carried out using PhyloBayes version 4.1. For both plastid and nuclear datasets, ten training datasets of 10,000 positions were used, and likelihoods were computed on ten test datasets of 2,000 positions.

### Data availability

The newly sequenced organelle genomes and their corresponding annotations are available online in the NCBI databases with accession numbers ranging from MK546602 to MK546611. The alignments used in this study as well as the resulting phylogenetic trees are available on figshare at the following address: https://doi.org/10.6084/m9.figshare.7680395.v2.

## Supporting information

All supplementary information

## Acknowledgments

We thank Zehra Çebi for growing up algal strains and for extraction of DNA, and Paul Simion and Rik Verdonck for critical reading of the manuscript. Sequencing was carried out by the Cologne Center for Genomics (CCG). Computations were performed on the supercomputers Mp2 and Ms2 from the Université de Sherbrooke, managed by Calcul Québec and Compute Canada. The operation of this supercomputer is funded by the Canada Foundation for Innovation (CFI), the ministère de l’Économie, de la science et de l’innovation du Québec (MESI), and the Fonds de recherche du Québec - Nature et technologies (FRQ-NT). This work was supported by the TULIP Laboratory of Excellence (ANR-10-LABX-41).

## Notes

### Competing Interest Statement

The authors have declared no competing interest.

## References

Archibald, John M. 2015. “Endosymbiosis and Eukaryotic Cell Evolution.” Current Biology 25 (19): R911–21. https://doi.org/10.1016/j.cub.2015.07.055.

Baurain, Denis, Henner Brinkmann, Jörn Petersen, Naiara Rodríguez-Ezpeleta, Alexandra Stechmann, Vincent Demoulin, Andrew J. Roger, Gertraud Burger, B. Franz Lang, and Hervé Philippe. 2010. “Phylogenomic Evidence for Separate Acquisition of Plastids in Cryptophytes, Haptophytes, and Stramenopiles.” Molecular Biology and Evolution 27 (7): 1698–1709. https://doi.org/10.1093/molbev/msq059.

Baurain, Denis, and Hervé Philippe. 2010. “Current Approaches to Phylogenomic Reconstruction.” In Evolutionary Genomics and Systems Biology, 17–41. Hoboken, NJ, USA: John Wiley & Sons, Inc. https://doi.org/10.1002/9780470570418.ch2.

Brown, Joseph W., and Ulf Sorhannus. 2010. “A Molecular Genetic Timescale for the Diversification of Autotrophic Stramenopiles (Ochrophyta): Substantive Underestimation of Putative Fossil Ages.” PLoS ONE 5 (9): 1–11. https://doi.org/10.1371/journal.pone.0012759.

Burki, Fabien, Kamran Shalchian-Tabrizi, Marianne Minge, Åsmund Skjæveland, Sergey I. Nikolaev, Kjetill S. Jakobsen, and Jan Pawlowski. 2007. “Phylogenomics Reshuffles the Eukaryotic Supergroups.” Edited by Geraldine Butler. PLoS ONE 2 (8): e790. https://doi.org/10.1371/journal.pone.0000790.

Delsuc, Frédéric, Georgia Tsagkogeorga, Nicolas Lartillot, and Hervé Philippe. 2008. “Additional Molecular Support for the New Chordate Phylogeny.” Genesis 46 (11): 592–604. https://doi.org/10.1002/dvg.20450.

Derelle, Romain, Purificación López-García, Hélène Timpano, and David Moreira. 2016. “A Phylogenomic Framework to Study the Diversity and Evolution of Stramenopiles (=Heterokonts).” Molecular Biology and Evolution 33 (11): 2890–98. https://doi.org/10.1093/molbev/msw168.

Dorrell, Richard G, Gillian Gile, Giselle McCallum, Raphaël Méheust, Eric P Bapteste, Christen M Klinger, Loraine Brillet-Guéguen, Katalina D Freeman, Daniel J Richter, and Chris Bowler. 2017. “Chimeric Origins of Ochrophytes and Haptophytes Revealed through an Ancient Plastid Proteome.” ELife 6 (May). https://doi.org/10.7554/eLife.23717.

Edgar, Robert C. 2010. “Search and Clustering Orders of Magnitude Faster than BLAST.” Bioinformatics 26 (19): 2460–61. https://doi.org/10.1093/bioinformatics/btq461.

Emms, David M, and Steven Kelly. 2015. “OrthoFinder: Solving Fundamental Biases in Whole Genome Comparisons Dramatically Improves Orthogroup Inference Accuracy.” Genome Biology 16 (1): 1–14. https://doi.org/10.1186/s13059-015-0721-2

Felsenstein, J. 1983. “Parsimony in Systematics: Biological and Statistical Issues.” Annual Review of Ecology and Systematics 14 (1): 313–33. https://doi.org/10.1146/annurev.es.14.110183.001525.

Felsenstein, Joseph. 1978. “Cases in Which Parsimony or Compatibility Methods Will Be Positively Misleading.” Systematic Zoology 27 (4): 401. https://doi.org/10.2307/2412923.

Felsenstein, Joseph. 1989. “PHYLIP - Phylogeny Inference Package - v3.2.” Cladistics. https://doi.org/10.1111/j.1096-0031.1989.tb00562.x.

Gatesy, John, and Mark S. Springer. 2014. “Phylogenetic Analysis at Deep Timescales: Unreliable Gene Trees, Bypassed Hidden Support, and the Coalescence/Concatalescence Conundrum.” Molecular Phylogenetics and Evolution 80 (1): 231–66. https://doi.org/10.1016/j.ympev.2014.08.013.

Gee, Henry. 2003. “Ending Incongruence.” Nature 425 (6960): 782–782. https://doi.org/10.1038/425782a.

Germot, Agnès, and Hervé Philippe. 1999. “Critical Analysis of Eukaryotic Phylogeny: A Case Study Based on the HSP70 Family.” The Journal of Eukaryotic Microbiology 46 (2): 116–24. https://doi.org/10.1111/j.1550-7408.1999.tb04594.x.

Graf, Louis, Eun Chan Yang, Kwi Young Han, Frithjof C. Küpper, Kylla M. Benes, Jason K. Oyadomari, Roger J.H. Herbert, et al. 2020. “Multigene Phylogeny, Morphological Observation and Re-Examination of the Literature Lead to the Description of the Phaeosacciophyceae Classis Nova and Four New Species of the Heterokontophyta SI Clade.” Protist 171 (6): 125781. https://doi.org/10.1016/j.protis.2020.125781.

Hahn, Matthew W., and Luay Nakhleh. 2016. “Irrational Exuberance for Resolved Species Trees.” Evolution 70 (1): 7–17. https://doi.org/10.1111/evo.12832.

Han, Kwi Young, Louis Graf, Carolina P. Reyes, Barbara Melkonian, Robert A. Andersen, Hwan Su Yoon, and Michael Melkonian. 2018. “A Re-Investigation of Sarcinochrysis Marina (Sarcinochrysidales, Pelagophyceae) from Its Type Locality and the Descriptions of Arachnochrysis, Pelagospilus, Sargassococcus and Sungminbooa Genera Nov.” Protist 169 (1): 79–106. https://doi.org/10.1016/j.protis.2017.12.004.

Heled, J., and A. J. Drummond. 2010. “Bayesian Inference of Species Trees from Multilocus Data.” Molecular Biology and Evolution 27 (3): 570–80. https://doi.org/10.1093/molbev/msp274.

Irisarri, Iker, Denis Baurain, Henner Brinkmann, Frédéric Delsuc, Jean Yves Sire, Alexander Kupfer, Jörn Petersen, et al. 2017. “Phylotranscriptomic Consolidation of the Jawed Vertebrate Timetree.” Nature Ecology and Evolution 1 (9): 1370–78. https://doi.org/10.1038/s41559-017-0240-5.

Kai, Atsushi, Yukie Yoshii, Takeshi Nakayama, and Isao Inouye. 2008. “Aurearenophyceae Classis Nova, a New Class of Heterokontophyta Based on a New Marine Unicellular Alga Aurearena Cruciata Gen. et Sp. Nov. Inhabiting Sandy Beaches.” Protist 159 (3): 435–57. https://doi.org/10.1016/J.PROTIS.2007.12.003.

Kalyaanamoorthy, Subha, Bui Quang Minh, Thomas K F Wong, Arndt von Haeseler, and Lars S Jermiin. 2017. “ModelFinder: Fast Model Selection for Accurate Phylogenetic Estimates.” Nature Methods 14 (6): 587–89. https://doi.org/10.1038/nmeth.4285.

Keeling, Patrick J., Fabien Burki, Heather M. Wilcox, Bassem Allam, Eric E. Allen, Linda a. Amaral-Zettler, E. Virginia Armbrust, et al. 2014. “The Marine Microbial Eukaryote Transcriptome Sequencing Project (MMETSP): Illuminating the Functional Diversity of Eukaryotic Life in the Oceans through Transcriptome Sequencing.” PLoS Biology 12 (6). https://doi.org/10.1371/journal.pbio.1001889.

Kolaczkowski, Bryan, and Joseph W. Thornton. 2004. “Performance of Maximum Parsimony and Likelihood Phylogenetics When Evolution Is Heterogeneous.” Nature 431 (7011): 980–84. https://doi.org/10.1038/nature02917.

Lartillot, Nicolas, Henner Brinkmann, and Hervé Philippe. 2007. “Suppression of Long-Branch Attraction Artefacts in the Animal Phylogeny Using a Site-Heterogeneous Model.” BMC Evolutionary Biology 7 (Suppl 1): S4. https://doi.org/10.1186/1471-2148-7-S1-S4.

Lartillot, Nicolas, and Hervé Philippe. 2004. “A Bayesian Mixture Model for Across-Site Heterogeneities in the Amino-Acid Replacement Process.” Molecular Biology and Evolution 21 (6): 1095–1109. https://doi.org/10.1093/molbev/msh112.

Lartillot, Nicolas, Nicolas Rodrigue, Daniel Stubbs, and Jacques Richer. 2013. “PhyloBayes MPI : Phylogenetic Reconstruction with Infinite Mixtures of Profiles in a Parallel Environment.” Systematic Biology 62 (4): 611–15. https://doi.org/10.1093/sysbio/syt022.

Le, Si Quang, Cuong Cao Dang, and Olivier Gascuel. 2012. “Modeling Protein Evolution with Several Amino Acid Replacement Matrices Depending on Site Rates.” Molecular Biology and Evolution 29 (10): 2921–36. https://doi.org/10.1093/molbev/mss112.

Lecointre, Guillaume, Hervé Philippe, Hoc Lanh Vân Lê, and Hervé Le Guyader. 1994. “How Many Nucleotides Are Required to Resolve a Phylogenetic Problem? The Use of a New Statistical Method Applicable to Available Sequences.” Molecular Phylogenetics and Evolution 3 (4): 292–309. https://doi.org/10.1006/mpev.1994.1037.

Lockhart, Peter, Phil Novis, Brook G. Milligan, Jamie Riden, Andrew Rambaut, and Tony Larkum. 2006. “Heterotachy and Tree Building: A Case Study with Plastids and Eubacteria.” Molecular Biology and Evolution 23 (1): 40–45. https://doi.org/10.1093/molbev/msj005.

Maddison, Wayne P. 1997. “Gene Trees in Species Trees.” Edited by John J. Wiens. Systematic Biology 46 (3): 523–36. https://doi.org/10.1093/sysbio/46.3.523.

Mirarab, Siavash, and Tandy Warnow. 2015. “ASTRAL-II: Coalescent-Based Species Tree Estimation with Many Hundreds of Taxa and Thousands of Genes.” Bioinformatics 31 (12): i44–52. https://doi.org/10.1093/bioinformatics/btv234.

Nadalin, Francesca, Francesco Vezzi, and Alberto Policriti. 2012. “GapFiller: A de Novo Assembly Approach to Fill the Gap within Paired Reads.” BMC Bioinformatics 13 (Suppl 14): S8. https://doi.org/10.1186/1471-2105-13-S14-S8.

Neiman, Maurine, and Douglas R Taylor. 2009. “The Causes of Mutation Accumulation in Mitochondrial Genomes.” Proceedings of the Royal Society B: Biological Sciences 276 (1660): 1201–9. https://doi.org/10.1098/rspb.2008.1758.

Nguyen, Lam-Tung T, Heiko A. Schmidt, Arndt von Haeseler, and Bui Quang Minh. 2015. “IQ-TREE: A Fast and Effective Stochastic Algorithm for Estimating Maximum-Likelihood Phylogenies.” Molecular Biology and Evolution 32 (1): 268–74. https://doi.org/10.1093/molbev/msu300.

Parks, Matthew B, Norman J Wickett, and Andrew J Alverson. 2018. “Signal, Uncertainty, and Conflict in Phylogenomic Data for a Diverse Lineage of Microbial Eukaryotes (Diatoms, Bacillariophyta).” Molecular Biology and Evolution 35 (1): 80–93. https://doi.org/10.1093/molbev/msx268.

Philippe, Hervé, Henner Brinkmann, Dennis V. Lavrov, D. Timothy J Littlewood, Michael Manuel, Gert Wörheide, and Denis Baurain. 2011. “Resolving Difficult Phylogenetic Questions: Why More Sequences Are Not Enough.” PLoS Biology 9 (3). https://doi.org/10.1371/journal.pbio.1000602.

Philippe, Hervé, Albert J. Poustka, Marta Chiodin, Katharina J. Hoff, Christophe Dessimoz, Bartlomiej Tomiczek, Philipp H. Schiffer, et al. 2019. “Mitigating Anticipated Effects of Systematic Errors Supports Sister-Group Relationship between Xenacoelomorpha and Ambulacraria.” Current Biology 29 (11): 1818–1826.e6. https://doi.org/10.1016/j.cub.2019.04.009.

Rodrigue, Nicolas, Hervé Philippe, and Nicolas Lartillot. 2010. “Mutation-Selection Models of Coding Sequence Evolution with Site-Heterogeneous Amino Acid Fitness Profiles.” Proceedings of the National Academy of Sciences of the United States of America 107 (10): 4629–34. https://doi.org/10.1073/pnas.0910915107.

Rogers, Scott O., and Arnold J. Bendich. 1985. “Extraction of DNA from Milligram Amounts of Fresh, Herbarium and Mummified Plant Tissues.” Plant Molecular Biology 5 (2): 69–76. https://doi.org/10.1007/BF00020088.

Roure, Béatrice, and Hervé Philippe. 2011. “Site-Specific Time Heterogeneity of the Substitution Process and Its Impact on Phylogenetic Inference.” BMC Evolutionary Biology 11 (1): 17. https://doi.org/10.1186/1471-2148-11-17.

Roure, Béatrice, Naiara Rodriguez-Ezpeleta, and Hervé Philippe. 2007. “SCaFoS: A Tool for Selection, Concatenation and Fusion of Sequences for Phylogenomics.” BMC Evolutionary Biology 7 (SUPPL. 1): 1–12. https://doi.org/10.1186/1471-2148-7-S1-S2.

Ševčíková, Tereza, Aleš Horák, Vladimír Klimeš, Veronika Zbránková, Elif Demir-Hilton, Sebastian Sudek, Jerry Jenkins, et al. 2015. “Updating Algal Evolutionary Relationships through Plastid Genome Sequencing: Did Alveolate Plastids Emerge through Endosymbiosis of an Ochrophyte?” Scientific Reports 5 (March): 1–12. https://doi.org/10.1038/srep10134.

Ševčíková, Tereza, Vladimír Klimeš, Veronika Zbránková, Hynek Strnad, Miluše Hroudová, Čestmír Vlček, and Marek Eliáš. 2016. “A Comparative Analysis of Mitochondrial Genomes in Eustigmatophyte Algae.” Genome Biology and Evolution 8 (3): 705–22. https://doi.org/10.1093/gbe/evw027.

Si Quang, Le, Olivier Gascuel, and Nicolas Lartillot. 2008. “Empirical Profile Mixture Models for Phylogenetic Reconstruction.” Bioinformatics 24 (20): 2317–23. https://doi.org/10.1093/bioinformatics/btn445.

Sibbald, Shannon J., and John M. Archibald. 2020. “Genomic Insights into Plastid Evolution.” Genome Biology and Evolution 12 (7): 978–90. https://doi.org/10.1093/gbe/evaa096.

Simion, Paul, Delsuc, Frédéric and Philippe, Hervé. To What Extent Current Limits of Phylogenomics Can Be Overcome?. Scornavacca, Celine; Delsuc, Frédéric; Galtier, Nicolas. Phylogenetics in the Genomic Era, No commercial publisher | Authors open access book, pp.2.1:1–2.1:34, 2020. hal-02535366

Simion, Paul, Hervé Philippe, Denis Baurain, Muriel Jager, Daniel J. D.J. Richter, Arnaud Di Franco, Béatrice Roure, et al. 2017. “A Large and Consistent Phylogenomic Dataset Supports Sponges as the Sister Group to All Other Animals.” Current Biology 27 (7): 958–67. https://doi.org/10.1016/j.cub.2017.02.031.

Simpson, Jared T, Kim Wong, Shaun D Jackman, Jacqueline E Schein, Steven J M Jones, and Inanç Birol. 2009. “ABySS: A Parallel Assembler for Short Read Sequence Data.” Genome Research 19 (6): 1117–23. https://doi.org/10.1101/gr.089532.108.

Springer, M. S., R. W. DeBry, C. Douady, H. M. Amrine, O. Madsen, W. W. de Jong, and M. J. Stanhope. 2001. “Mitochondrial versus Nuclear Gene Sequences in Deep- Level Mammalian Phylogeny Reconstruction.” Molecular Biology and Evolution, no. 18: 132–43.

Stamatakis, Alexandros. 2014. “RAxML Version 8: A Tool for Phylogenetic Analysis and Post-Analysis of Large Phylogenies.” Bioinformatics 30 (9): 1312–13. https://doi.org/10.1093/bioinformatics/btu033.

Whitfield, James B., and Peter J. Lockhart. 2007. “Deciphering Ancient Rapid Radiations.” Trends in Ecology and Evolution 22 (5): 258–65. https://doi.org/10.1016/j.tree.2007.01.012.

Yang, Eun Chan, Ga Hun Boo, Hee Jeong Kim, Sung Mi Cho, Sung Min Boo, Robert A. Andersen, and Hwan Su Yoon. 2012. “Supermatrix Data Highlight the Phylogenetic Relationships of Photosynthetic Stramenopiles.” Protist 163 (2): 217–31. https://doi.org/10.1016/j.protis.2011.08.001.

Zhang, Chao, Maryam Rabiee, Erfan Sayyari, and Siavash Mirarab. 2018. “ASTRAL-III: Polynomial Time Species Tree Reconstruction from Partially Resolved Gene Trees.” BMC Bioinformatics 19 (S6): 153. https://doi.org/10.1186/s12859-018-2129-y.

Zwickl, Derrick J., and David M. Hillis. 2002. “Increased Taxon Sampling Greatly Reduces Phylogenetic Error.” Edited by Keith Crandall. Systematic Biology 51 (4): 588–98. https://doi.org/10.1080/10635150290102339.

