## Supplementary material for "Lower statistical support with larger datasets: insights from the Ochrophyta radiation": All supplementary information

#### Methods

##### 1. Creation of the plastid dataset

We retrieved the protein annotations for 75 selected plastid genomes of Rhodophyta, Cryptophyta, Haptophyceae and Ochrophyta from the NCBI RefSeq database (https://www.ncbi.nlm.nih.gov/) (Supplementary Table 8). We used OrthoFinder (Emms and Kelly 2015) with a BLASTP E-value threshold of 1e-5 and an MCL inflation parameter of 1.5 to produce orthogroups (OGs). We filtered the 504 resulting OGs to retain those (108) containing ≥20 species (of which ≥1 Rhodophyta, ≥1 Stramenopiles, and either ≥1 Cryptophyta or ≥1 Haptophyceae). We first aligned the selected OGs with MAFFT (L-INS-i algorithm, 5000 iterations) (Katoh and Standley 2013), then enriched them by adding more species from genomic data (such as the five new species sequenced in this study) with Forty-Two (https://metacpan.org/release/Bio-MUST-Apps-FortyTwo). We checked for possible paralogy using methods that are described in the section about the construction of the nuclear dataset (see below) and found only one dubious OG, from which we manually removed four paralogous sequences. We further discarded 9 additional OGs with <30 species. Finally, to select unambiguously aligned positions, we applied a loose BMGE (Criscuolo et al. 2010) filter (entropy cutoff of 0.6 and gap cutoff of 0.4) on each aligned OG.

##### 2. Creation of the mitochondrial dataset

As for the plastid, we retrieved all the protein annotations available for stramenopiles mitochondrial genomes from the NCBI website (Supplementary Table 9). To this first set, we added the annotations of the five new species generated in this study, as well as some identified from genomic scaffolds of Labyrinthulomycetes and Xanthophyceae species presenting a high similarity to mitochondrial genomes, using MFannot server (Beck and Lang 2010; MFannot, organelle genome annotation websever; http://megasun.bch.umontreal.ca/cgi-bin/mfannot/mfannotInterface.pl). We chose to integrate all annotations before delineating OGs because it can be more difficult to retrieve orthologs for fast evolving mitochondrial sequences with Forty-Two. We generated OGs using the same protocol as with the plastid dataset and retained the 33 OGs containing ≥66% of the species of the dataset. We aligned the clusters with MAFFT (L-INS-i algorithm, 5000 iterations) and manually fixed frameshift errors for some sequences of *Fragilariopsis* and *Aurantiochytrium* in the resulting alignments. We also split the sequence of *Phaeodactylum* fusion protein nad9-rps14 and added each half to its respective alignment. Finally, we removed one OG showing a too ambiguous alignment and applied BMGE on each aligned OG as above.

##### 3. Generation and selection of orthogroups for the nuclear dataset

We retrieved the complete proteomes for 53 species across Stramenopiles, Alveolata and Rhizaria from the NCBI website (Supplementary Table 10). We performed pairwise similarity searches between all proteomes using USEARCH (Edgar 2010) with a minimal value of 1e-5 (instead of BLAST to speed up computations). We then generated OGs with OrthoFinder, as for the plastid and mitochondrial datasets. Out of a total of 212,849 OGs, we retained only those containing ≥15 species, with at least one species from each of the three clades (Stramenopiles, Alveolata and Rhizaria) and ≤600 sequences to avoid retrieving large protein families. These filters left us with 3,063 OGs.

At this stage, we observed that some OGs contained highly divergent sequences (possibly non-homologous), dragged into the clusters by a single similarity link. To address this issue, we removed the sequences that were not similar to a minimum percentage of the other sequences in each OG (BLAST E-value threshold of 1e-10). We proceeded in two steps, first removing the sequences matching <30% of the other sequences, then 50%. These two steps removed 17,734 and 6,335 sequences, respectively. Finally, we filtered the OGs anew, so as to retain only those with ≥15 species, hence reducing their number to 2,892.

To classify OGs among those close to true orthogroups and those corresponding to more complex multigene families, we used an automated phylogenetic analysis of single-gene trees (Simion et al., 2017). Briefly, we first aligned OGs with MAFFT v7 (L-INS-I algorithm, 5000 iterations) (Katoh and Stanley 2013), then filtered out columns with <5% of amino acid residues and sequences with <50 parsimony informative positions. We then inferred trees with RAxML v8 (Stamatakis 2014) using the LG+F+Γ4 model (Le and Gascuel 2008), and ran a custom script aimed at detecting cases of ancient paralogy. As in Simion et al. (2017), we computed how many of seven predefined clades (Rhizaria, Ciliophora, Myzozoa, Oomycetes, Labyrinthulomycetes, *Blastocystis* and Ochrophyta) were affected by out-paralogy (i.e., at least two clans containing sequences exclusively from a given clade). This allowed us to separate the OGs containing ≤3 out-paralogs (1904) from the other, more complex OGs (988) having more out-paralogs.

To split the OGs showing too many cases of out-paralogy (which may correspond to an ancient gene duplication), we used the software root-max-div (Simion et al. 2017). This program searches for the branch maximizing the taxonomic diversity on both sides of the bipartition and splits the tree on the branch if (i) the number of sequences on each side satisfies a minimal threshold (two first parameters), (ii) the number of common species on both sides is above a minimal threshold (third parameter) and (iii) the branch length is among the top percentile of the tree branches (fourth parameter). We applied four different parameter sets in the following order (30-30-0-5, 30-10-0-5, 10-10-0-5, and 40-10-0-20), retrieving the two sub-alignment files of the first successful parameter set for each OG. We repeated the whole procedure until no more gene tree could be cut. Finally, we filtered the split OGs again, to retain only those with ≥15 species, thereby reducing their number to 336.

Finally, we pooled the two groups of OGs, yielding a total of 2,240 OGs, of which we reduced the redundancy with a custom script targeting highly similar subsequences of the same organism inside individual OGs, as in Simion et al. (2017). This step removed 5,573 sequences.

#### 4. Assessment of transcriptome quality

Before improving the taxon sampling of our nucleus dataset using a combination of transcriptomic and genomic data, we evaluated the contamination level of the available transcriptomes. They consisted in assemblies from MMETSP (Keeling 2014), TSA retrieved from the NCBI website, and SRA raw reads also retrieved from the NCBI website, which were assembled using Trinity v2.6 (Haas 2013) with the trimmomatic and jaccard clip options (--jaccard_clip --trimmomatic). A set of 80 highly expressed gene alignments (ribosomal proteins), on which a large diversity of eukaryotic sequences are regularly added and manually curated, was used as a reference. To estimate the contamination level, we took advantage of Forty-Two and its taxonomic filters to add back previously incorporated organisms to this dataset. Forty-Two was designed to search transcriptomes for orthologous sequences and add them to existing alignments. At this last step, it can verify if the added orthologous sequence satisfies a user-defined positive and/or negative taxonomic filter (i.e., belonging to Stramenopiles or not belonging to Xanthophyceae). Here, we checked that added sequences indeed matched an organism of the same genus in the alignment, which allowed us to distinguish between orthologous sequences genuinely belonging to the transcriptome from orthologous sequences belonging to contaminants. We considered a transcriptome to be clean when we found <5 contaminant sequences over the 80 alignments. For each contaminant sequence, we further retrieved the most closely related organism in the alignment, so as to design optimal taxonomic filters for contaminated transcriptomes. Overall, this approach allowed us to exploit taxonomically interesting transcriptomes that were contaminated without adding contaminated sequences in our OGs.

#### 5. *Vaucheria litorea* transcriptome decontamination

Whereas *Vaucheria litorea* was one of the only Xanthophyceae for which a large amount of data was available, we observed that its transcriptome was contaminated by a large array of organisms. Because of its isolated phylogenetic position in our current sample of the eukaryotic diversity, combined to a relatively fast evolutionary rate, it was difficult to only rely on sequence similarity for decontamination. Thus, to tackle this issue, we implemented a strategy based on k-mer distributions (Teeling et al. 2004) to identify and remove the largest part of *Vaucheria* contaminant sequences. Briefly, we assembled two sets of *Vaucheria* sequences (i.e., genuine and contaminant), for which we computed the frequencies of all possible 6-nt k-mers. Then, the k-mer composition of each transcript was compared against those reference distributions using an Euclidean distance, and we discarded the transcripts closer to the contaminant than the genuine sequences. To define these two sets of reference, we used eukaryotic orthologs (594 nuclear genes from 370 species covering the diversity of eukaryotes) from a non-published study, in which we added *Vaucheria* transcripts with Forty-Two. (We used these orthologs instead of the 80 ribosomal proteins to maximize the number and the variety of sequences in the reference sets.) Then, we inferred the single-gene phylogeny of all orthologs using RAxML (LG+F+Γ4 model) and retrieved the taxonomy of the sister clan of each *Vaucheria* sequence. We considered transcripts added close to Ochrophyta species as genuine reference sequences, whereas the other transcripts were pooled separately as different sources of contaminants. Those were mainly representatives of Labyrinthulomycetes, Discosea and Viridiplantae. To identify contaminants in the full *Vaucheria* transcriptome, we first tested the transcripts against Labyrinthulomycetes contaminated sequences then to Discosea sequences, and finally to Viridiplantae sequences. We verified the effectiveness of our approach with the protocol described in section 4, which confirmed that the majority of the contaminants had been properly removed after using the Labyrinthulomycetes and Discosea references. Thus, we used the resulting transcriptome in our dataset construction without excluding the sequences that would have been removed after testing against Viridiplantae.

#### 6. Enrichment of OGs to increase taxonomic diversity

Starting from OGs generated at section 3, we improved our taxonomic sampling by adding sequences from genomic and transcriptomic data with Forty-Two. We worked in two consecutive steps, first adding non-contaminated transcriptomic data (127 species) (see section 4. Assessment of transcriptomes quality) and genomic data (13 species), and then contaminated transcriptomic data (54 species). As explained in section 4, we took advantage of the presence of contaminants relatives or sequenced organisms relatives in our OGs to design custom taxonomic filters for the added sequences (see YAML files in Supplementary Archive). After each run of Forty-Two, we ran the custom script described at the end of section 3 to reduce the number of redundant sequences per species. Finally, we filtered the enriched OGs to retain only those with ≥5 species of Rhizaria, ≥10 species of Alveolata, ≥10 non-photosynthetic Stramenopiles and ≥20 photosynthetic Stramenopiles (Ochrophyta), leaving us with 1330 enriched OGs and a total of 244 species.

#### 7. Targeted decontamination

Some transcriptomes showed evidence of contamination by organisms outside of SAR, which cannot be handled by the taxonomic filters of Forty-Two (due to the lack of related sequences in our OGs). To remove these contaminant sequences from our OGs, we used a custom script to BLAST each sequence against two reference databases of wanted (i.e., SAR species) and unwanted (i.e., contaminant species) proteomes. We then discarded the sequences better matching the unwanted database over the wanted database. Contamination sources detected at this step were as various as red algae (70 sequences), green algae (302 sequences), green plants (297 sequences), animals (247 sequences), fungi (12 sequences) or alpha-proteobacteria (3481 sequences, probably containing some genuine sequences of mitochondrial origin).

#### 8. Detection and elimination of remaining paralogs

After targeted removal of contaminant sequences, we realigned each OG with MAFFT and searched for possibly remaining paralogs that could not have been handled by our tree splitting step and/or that appeared during OG enrichment. We inferred single-gene trees as before (RAxML, LG+F+Γ4 model) and used them to split alignments with the previously described script, but using new values of the four parameters (50-50-50-10). We also split trees for recent paralogs specific to one clade (at least 150 species on one side, 5 on the other side, 3 in common, and branch length among the top-10%, 150-5-3-10), discarding the smallest sub-alignment file. This step mainly allowed us to detect and remove potential nucleomorph sequences. The latter sequences were more specifically targeted by isolating Rhizaria sequences, splitting them with a 10-1-1-5 parameter set and discarding the smallest group of rhizarian sequences. Finally, we reassessed the presence of ancient paralogs between the 7 clades, as described in section 3 but with a small modification. We inferred single gene trees without the sequences with <100 parsimony informative positions, as misplacement of these fragments could inflate the separation of monophyletic clades (Di Franco et al. 2019). We kept the clusters with less than 16 out-paralogs and filtered them for ≥93 species, leaving us with a total of 1,119 alignments of orthologous sequences.

#### 9. Branch length decontamination and filtering

To remove remaining outlier sequences of various origins (contaminants, paralogs or xenologs), we used the same protocol as in Simion et al. (2017). Instead of using a block-oriented BMGE filter on our OGs as above, we applied HmmCleaner v1.8 (Di Franco et al. 2019) with default parameters and filtered columns with <5% of amino acid residues, in order to keep as much signal as possible for inferring single-gene trees. We created a supermatrix with those OGs using SCaFoS (Roure et al. 2007), picking the longest sequence per OTU. On one side, we inferred the supermatrix tree with RAxML and LG+F+Γ4 model to serve as the reference tree. On the other side, we inferred the branch lengths of each individual OG while constraining the tree topology to the supermatrix topology. Finally, we compared the terminal branch lengths of single-gene trees to those of the supermatrix tree for each OTU. We performed this procedure twice, removing sequences that were x times longer in a single-gene tree compared to the supermatrix tree. We first removed 423 sequences using a branch length ratio of x=7 and then 465 sequences using a ratio of x=5. Moreover, we discarded 53 OGs in which too many branch lengths were incoherent with those of the supermatrix tree. To identify these OGs, we computed a branch length $R^2$ for each OG with respect to the supermatrix tree, and set the elimination threshold to a $R^2$ value below the mean value of the $R^2$ distribution minus 1.96 times the standard deviation, assuming a normal distribution of $R^2$ values.

In the last step of our nuclear dataset construction, we reduced our taxon sampling to 185 SAR OTUs by either removing too closely related species or building chimeras with SCaFoS, depending on their completeness. We applied BMGE as above, followed by HmmCleaner with default parameters on each OG. We retained the 797 OGs having ≤62 missing OTUs using SCaFoS, and subsequently removed Rhizaria and Alveolata.

#### 10. DNA extraction via CTAB method reagents (modified protocol)

reagents

- TE buffer: 10 mM Tris-HCl (pH 8), 1mM EDTA, sterile bidistilled water
- 5% SDS solution
- proteinase K solution: 10 mg/ ml
- RNAse: 10 mg/ml
- 5M NaCl

- CTAB buffer (pre-warmed at 65°C): 1.4 M NaCl, 0.1 M Tris-HCl (pH 8), 25 mM EDTA, 2% CTAB (w/v), sterile bidistilled water
- 24:1 chloroform : isoamyl alcohol
- Isopropanol/2-propanol (pre-cooled at -20°C)
- 80% ethanol (pre-cooled at -20°C)

protocol

1. harvest algae & wash pellet several times with respective medium (centrifugation at lowest-possible speed)
2. homogenize pellet (≈ 1 ml) with liquid nitrogen
3. transfer homogenized powder very quickly into a sterile falcon tube containing a mixture of 900 µl TE buffer, 700 µl 5% SDS and 25 µl proteinase K solution & vortex immediately
4. incubate in water bath at 60°C, 20‘ (vortex occasionally)
5. add 500 µl 5 M NaCl, 25 µl RNAse A and 5 ml CTAB buffer (pre-warmed at 65°C) & vortex
6. incubate in water bath at 60°C, 10‘ (vortex occasionally)
7. centrifuge the sample (3,000 g, 15‘) & transfer supernatant into a new sterile falcon tube
8. add equal volume of 24:1 chloroform : isoamyl alcohol & vortex rigorously
9. incubate at RT, 10’
10. separate phases by centrifugation (3,000 g, 15’)
11. transfer & portion the upper, aqueous phase carefully into sterile 1.5 ml eppendorf tubes (if the aqueous phase is not clear, the chloroform-isoamyl alcohol extraction has to be repeated from step 8)
12. maintain DNA extracts on ice & perpetuate this condition
13. add 2/3 volume of isopropanol (pre-cooled at -20°C) & vortex
14. incubate at -20°C, at least 1 h (or overnight)
15. centrifuge (17,000 g, 15’, 4°C) & discard the supernatant carefully
16. wash pellet with 1 ml 80% ethanol (pre-cooled at -20°C) & vortex
17. centrifuge (17,000 g, 5’, 4°C) & discard the supernatant carefully
18. repeat the washing step
19. air-dry the pellet & dissolve the pellet in 50 - 200 µl TE buffer
20. store at -20°C

### References


Criscuolo A, Gribaldo S. 2010. BMGE (Block Mapping and Gathering with Entropy): a new software for selection of phylogenetic informative regions from multiple sequence alignments. BMC Evol. Biol. 10:210.

Di Franco A, Poujol R, Baurain D, Philippe H. 2019. Evaluating the usefulness of alignment filtering methods to reduce the impact of errors on evolutionary inferences. BMC Evol. Biol. 19:21.

Edgar RC. 2010. Search and clustering orders of magnitude faster than BLAST. 26:2460–2461.

Emms DM, Kelly S. 2015. OrthoFinder: solving fundamental biases in whole genome comparisons dramatically improves orthogroup inference accuracy. Genome Biol. 16:1–14.

Haas BJ, Papanicolaou A, Yassour M, Grabherr M, Blood PD, Bowden J, Couger MB, Eccles D, Li B, Lieber M, et al. 2013. De novo transcript sequence reconstruction from RNA-seq using the Trinity platform for reference generation and analysis. Nat. Protoc. 8:1494–1512.

Katoh K, Standley DM. 2013. MAFFT multiple sequence alignment software version 7: improvements in performance and usability. Mol Biol Evol 30:772–780.

Keeling PJ, Burki F, Wilcox HM, Allam B, Allen EE, Amaral-Zettler L a., Armbrust EV, Archibald JM, Bharti AK, Bell CJ, et al. 2014. The Marine Microbial Eukaryote Transcriptome Sequencing Project (MMETSP): Illuminating the Functional Diversity of Eukaryotic Life in the Oceans through Transcriptome Sequencing. PLoS Biol. 12.

Le SQ, Gascuel O. 2008. An Improved General Amino Acid Replacement Matrix. Mol. Biol. Evol. 25:1307–1320.

Roure B, Rodriguez-Ezpeleta N, Philippe H. 2007. SCaFoS: A tool for selection, concatenation and fusion of sequences for phylogenomics. BMC Evol. Biol. 7:1–12.

Simion P, Philippe H, Baurain D, Jager M, Richter DJDJ, Di Franco A, Roure B, Satoh N, Quéinnec É, Ereskovsky A, et al. 2017. A Large and Consistent Phylogenomic Dataset Supports Sponges as the Sister Group to All Other Animals. Curr. Biol. 27:958–967.

Stamatakis A. 2014. RAxML version 8: A tool for phylogenetic analysis and post-analysis of large phylogenies. Bioinformatics 30:1312–1313.

Teeling H, Meyerdierks A, Bauer M, Amann R, Glockner FO. 2004. Application of tetranucleotide frequencies for the assignment of genomic fragments. Environ. Microbiol. 6:938–947.

### Supplementary Table 1. Bootstrap support of high-level ochrophyte clades for 23 species datasets with varying taxon sampling under LG4X model

| | Plastid | | | | | | | | Nucleus | | | | | | | |
|---|---|---|---|---|---|---|---|---|---|---|---|---|---|---|---|---|
| Groupings | All | distant | E | SSC | Ping | PXR | PD | BB | All | distant | E | SSC | Ping | PXR | PD | BB |
| BB+E+PD+PXR | | | . | | | . | . | . | | | . | | **60** | . | . | . |
| BB+E+PD+PXR+SSC | | | . | . | | . | . | . | | 76 | . | . | | . | . | . |
| BB+PD | 54 | 96 | 61 | 46 | 64 | 88 | . | . | 51 | 81 | 70 | | 100 | 59 | . | . |
| BB+PD+Ping | | | | | . | | . | . | | | | **98** | . | **81** | . | . |
| BB+PD+Ping+PXR | | **54** | | | . | . | . | . | | | | *97* | . | . | . | . |
| BB+PD+Ping+PXR+SSC | | | | . | . | . | . | . | 33 | | | . | . | . | . | . |
| BB+PD+Ping+SSC | | | | . | . | | . | . | 36 | | 51 | . | . | | . | . |
| BB+PD+PXR | | **80** | | | | . | . | . | | | | | | . | . | . |
| BB+Ping | | | | | . | | | . | | | | **67** | . | | **96** | . |
| **E+Ping** | | | . | 78 | . | | | | | | . | | . | | | |
| E+Ping+PXR | | | . | 97 | . | . | | | | | . | | . | . | | |
| E+Ping+PXR+SSC | 99 | | . | . | . | . | 100 | 100 | | | . | . | . | . | | |
| E+Ping+SSC | 93 | | . | . | . | 100 | 68 | 84 | | | . | . | . | | | |
| **E+PXR** | | | . | | | . | | | | | . | | **59** | . | | |
| E+PXR+SSC | | | . | . | 99 | . | | | | 71 | . | . | | . | **44** | |
| **E+SSC** | 100 | 100 | . | . | 100 | 100 | 100 | 100 | | 71 | . | . | | **60** | **47** | |
| PD+Ping+PXR+SSC | | | | . | . | . | . | | | | | . | . | . | . | *47* |
| PD+Ping+SSC | | | | . | . | | . | | | | | . | . | | . | 47 |
| Ping+PXR+SSC | | | 98 | . | . | . | | | | | | . | . | . | | |
| **Ping+SSC** | | | 100 | . | . | | | | 35 | | 51 | . | . | | | 44 |

Rows correspond to the observed high-level groupings and columns to major clades that were left out from the taxon sampling (All means that all species are considered). Dots (.) indicate groupings not testable with the corresponding taxon sampling of the column, italics indicate groupings that are compatible, but not directly comparable, to the corresponding grouping formed when all the species are considered, boldface indicates groupings that are not observed when all the species are considered. Abbreviations are as in Fig. 2, and Out means use of a distant outgroup (i.e., removal of the close outgroup).

### Supplementary Table 2. Comparison of evolutionary models

| Compartment | Model | Likelihood (ln L) | Degrees of freedom | AIC | AICc | BIC |
|---|---|---|---|---|---|---|
| Plastid | LG4X | -817,778.87 | 129 | 1,635,815.74 | 1,635,817.30 | 1,636,845.77 |
| Plastid | LG+C20+G | -811,660.60 | 124 | 1,623,569.20 | 1,623,570.64 | 1,624,559.30 |
| Nucleus | LG4X | -12,949,987.47 | 251 | 25,900,476.94 | 25,900,477.55 | 25,903,049.84 |
| Nucleus | LG+C20+G | -12,716,958.52 | 246 | 25,434,409.04 | 25,434,409.62 | 25,436,930.69 |

### Supplementary Table 3. Support of high-level ochrophyte clades comparison between bootstrap replicates of full matrix and Jackknife replicates of 50.000 bp

| | Nucleus bootstrap 210000 | | | | | | | | Nucleus JK50000 50 rep | | | | | | | |
|---|---|---|---|---|---|---|---|---|---|---|---|---|---|---|---|---|
| Groupings | All | Out | E | SSC | Ping | PXR | PD | BB | All | Out | E | SSC | Ping | PXR | PD | BB |
| BB+E+PD+PXR | | **78** | . | | | . | . | . | | | . | | | . | . | . |
| BB+E+PD+PXR+ SSC | | 100 | . | . | | . | . | . | | 78 | . | . | | . | . | . |
| BB+PD | 68 | 100 | 97 | | 100 | | . | . | 57 | 70 | 60 | | 100 | | . | . |
| BB+PD+Ping | | | | 95 | . | 88 | . | . | | | | 74 | . | 76 | . | . |
| BB+Ping | | | | 95 | . | 88 | 83 | . | | | | 66 | . | 72 | 75 | . |
| E+Ping+PXR+ SSC | 67 | | . | . | . | . | | 100 | 56 | | . | . | . | . | | 54 |
| **E+PXR** | 81 | 82 | . | 96 | | . | 91 | 100 | 76 | 72 | . | 80 | | . | 77 | 94 |
| E+PXR+SSC | | | . | . | *83* | . | 83 | | | **48** | . | . | *80* | . | 65 | |
| **E+SSC** | | | . | . | 56 | 91 | | | | | . | . | 54 | 84 | | |
| Ping+PXR+SSC | | | *96* | . | . | . | | | | | *60* | . | . | . | | |
| **Ping+SSC** | 68 | | 97 | . | . | | | 100 | 56 | | 64 | . | . | | | 62 |

See legend of Supplementary Table 1 for details

### Supplementary Table 4. Bootstrap support of high-level ochrophyte clades with varying taxon sampling under C20+LG model

| | Plastid | | | | | | | | Nucleus | | | | | | | |
|---|---|---|---|---|---|---|---|---|---|---|---|---|---|---|---|---|
| Groupings | All | Out | E | SSC | Ping | PXR | PD | BB | All | Out | E | SSC | Ping | PXR | PD | BB |
| BB+E+PD+PXR | | | . | | | . | . | . | | **35** | . | | | . | . | . |
| BB+E+PD+PXR+SSC | | | . | . | | . | . | . | | **74** | . | . | | . | . | . |
| BB+PD | 99 | 100 | 99 | 94 | 100 | 94 | . | . | 100 | 100 | 100 | 99 | 100 | 99 | . | . |
| BB+PD+Ping | | | | | . | *72* | . | . | | | | | . | | . | . |
| BB+PD+Ping+PXR | 74 | 98 | | 86 | . | . | . | . | | | | | . | . | . | . |
| BB+PD+PXR | | **95** | | | *94* | . | . | . | | | | | | . | . | . |
| BB+Ping | | | | | . | | **70** | . | | | | | . | | | . |
| BB+Ping+PXR | | | | | . | . | *82* | . | | | | | . | . | | . |
| E+Ping+PXR | | | . | | . | . | | | | | . | *95* | . | . | | |
| E+Ping+PXR+SSC | | | . | . | . | . | | | 99 | | . | . | . | . | 51 | 94 |
| E+Ping+SSC | | | . | . | . | | | | | | . | . | . | *81* | | |
| **E+PXR** | | | . | | | . | | | 100 | 84 | . | 100 | 52 | . | 96 | 100 |
| E+PXR+SSC | | | . | . | | . | | | | | . | . | *100* | . | | |
| **E+SSC** | 100 | 100 | . | . | 100 | 100 | 100 | 100 | | | . | . | | **92** | | |
| PD+Ping+PXR | | | | | . | . | . | *87* | | | | | . | . | . | |
| **Ping+PXR** | 64 | | | 62 | . | . | | 47 | | | | | . | . | | |
| Ping+PXR+SSC | | | **64** | . | . | . | | | | | *100* | . | . | | | |
| **Ping+SSC** | | | **67** | . | . | | | | 99 | | 100 | . | . | | 51 | 94 |

See legend of Supplementary Table 1 for details

### Supplementary Table 5. Statistical support of high-level clades for the 23-species fusion (nu+cp) dataset with varying taxon sampling under LG4X and CAT+Γ4 models

| | LG4X | | | | | | | CAT (jack 80000) | | | | | | |
|---|---|---|---|---|---|---|---|---|---|---|---|---|---|---|
| Groupings | All | E | SSC | Ping | PXR | PD | BB | All | E | SSC | Ping | PXR | PD | BB |
| BB+PD | | **86** | **59** | *100* | **74** | . | . | 100 | 98 | 99 | 100 | 99 | . | . |
| BB+PD+Ping | 65 | | 85 | . | 99 | . | . | 100 | 88 | 99 | . | 79 | . | . |
| BB+PD+Ping+ PXR | | | **64** | . | . | . | . | | | | . | . | . | . |
| BB+PD+Ping+ SSC | | **63** | . | . | | . | . | | | . | . | | . | . |
| BB+Ping | 46 | | | . | | 100 | . | | | | . | | *99* | . |
| **E+PXR** | | . | | | . | | | | . | *100* | | . | | |
| E+PXR+SSC | 86 | . | . | 94 | . | 100 | 67 | 90 | . | . | 96 | . | 96 | 86 |
| **E+SSC** | 100 | . | . | 100 | 100 | 100 | 100 | 99 | . | . | 100 | 100 | 100 | 98 |
| PD+Ping | | | | . | | . | *100* | | | | . | | . | *100* |
| **Ping+SSC** | | **64** | . | . | | | | | | . | . | | | |
| PXR+SSC | | | . | | . | | | | *70* | . | | . | | |

See legend of Supplementary Table 1 for details

Supplementary Table 6. Jackknife support of high-level clades for the 23-species fusion (nu+cp) matrix under CAT model over varying replicate size

| | CAT (jackknife 20,000 positions) | | | | | | | | CAT (jackknife 50,000 positions) | | | | | | | | CAT (jackknife 80,000 positions) | | | | | | | |
|---|---|---|---|---|---|---|---|---|---|---|---|---|---|---|---|---|---|---|---|---|---|---|---|---|
| Groupings | All | distant | E | SSC | Ping | PXR | PD | BB | All | distant | E | SSC | Ping | PXR | PD | BB | All | distant | E | SSC | Ping | PXR | PD | BB |
| BB+PD | 86 | 85 | 91 | 81 | 99 | 79 | . | . | 94 | 98 | 98 | 99 | 100 | 96 | . | . | 100 | 100 | 98 | 99 | 100 | 99 | . | . |
| BB+PD+ Ping | 72 | 54 | 56 | 82 | . | 60 | . | . | 95 | 90 | 71 | 92 | . | 40 | . | . | 100 | 96 | 88 | 99 | . | 79 | . | . |
| BB+PD+ Ping+SSC | | | **38** | . | . | | . | . | | | | . | . | | . | . | | | | . | . | | . | . |
| BB+Ping | | | | | . | | *65* | . | | | | | . | | *89* | . | | | | | . | | *99* | . |
| **E+PXR** | | | . | *81* | | . | | | | | . | *94* | | . | | | | | . | *100* | | . | | |
| E+PXR+ SSC | 50 | 67 | . | . | 68 | . | 49 | 45 | 85 | 87 | . | . | 88 | . | 71 | 71 | 90 | 98 | . | . | 96 | . | 96 | 86 |
| **E+SSC** | 77 | 78 | . | . | 93 | 94 | 83 | 76 | 98 | 99 | . | . | 98 | 100 | 96 | 92 | 99 | 99 | . | . | 100 | 100 | 100 | 98 |
| PD+Ping | | | | | . | | . | *73* | | | | | . | | . | *93* | | | | | . | | . | *100* |
| PXR+SSC | | | | . | | . | | | | | *55* | . | | . | | | | | *70* | . | | . | | |

See legend of Supplementary Table 1 for details. Average bootstrap support is increasing with replicate size from 72.38 at 20.000 positions to 88.04 at 50.000 positions and finally 95.73 at 80.000 positions.

### Supplementary Table 7. Species matches between compartments for common taxon sampling analyses

| Mitochondrion | Plastid | Nucleus |
|---|---|---|
| *Phytophthora sojae* | *Guillardia theta* | *Phytophthora parasitica* |
| *Aureococcus anophagefferens* | *Aureococcus anophagefferens* | *Aureococcus anophagefferens* |
| *Sarcinochrysidales sp._CCMP2135* | *Aureoumbra lagunensis* | *Aureoumbra lagunensis_CCMP1510* |
| *Chromulina chionophila* | *Chromulina chionophila_CCAP9099* | *Chromulina nebulosa_UTEXLB2642* |
| *Berkeleya fennica* | *Coscinodiscus radiatus* | *Coscinodiscus wailesii_CCMP2513* |
| *Ectocarpus siliculosus* | *Ectocarpus siliculosus* | *Ectocarpus siliculosus* |
| *Florenciella parvula* | *Florenciella parvula_RCC446* | *Florenciella parvula_CCMP2471* |
| *Heterosigma akashiwo* | *Heterosigma akashiwo* | *Heterosigma akashiwo_CCMP2393* |
| *Nannochloropsis gaditana* | *Nannochloropsis gaditana* | *Nannochloropsis gaditana* |
| *Nannochloropsis limnetica* | *Nannochloropsis limnetica* | *Nannochloropsis limnetica* |
| *Nannochloropsis oceanica* | *Nannochloropsis oceanica* | *Nannochloropsis oceanica* |
| *Ochromonas danica* | *Ochromonas sp._CCMP1393* | *Ochromonas sp._CCMP1393* |
| *Asterionella formosa* | *Leptocylindrus danicus* | *Leptocylindrus danicus_B650* |
| *Phaeomonas parva* | *Phaeomonas parva_CCMP2877* | *Phaeomonas parva_CCMP2877* |
| *Pseudopedinella elastica* | *Pseudopedinella elastica_SAGB4388* | *Pseudopedinella elastica_CCMP716* |
| *Phaeodactylum tricornutum* | *Rhizosolenia imbricata* | *Rhizosolenia setigera_CCMP1694* |
| *Saccharina japonica* | *Saccharina japonica* | *Saccharina angustata* |
| *Sargassum thunbergii* | *Sargassum thunbergii* | *Sargassum thunbergii* |
| *Synura peterssenii* | *Synura peterssenii_CCAC0052* | *Mallomonas sp._CCMP3275* |
| *Trachydiscus minutus* | *Trachydiscus minutus* | *Eustigmatos cf._polyphem* |
| *Triparma laevis* | *Triparma laevis* | *Bolidomonas pacifica_CCMP1866* |
| *Heterococcus sp._DN1* | *Vaucheria litorea* | *Vaucheria litorea* |
| - | uncultured *Pelagomonas* | *Pelagomonas calceolata_CCMP1756* |

### Supplementary Table 8. Genomes used to construct the plastid dataset

| Species |
|---|
| Ahnfeltia plicata |
| Apophlaea sinclairii |
| Asparagopsis taxiformis |
| Asterionella formosa |
| Asterionellopsis glacialis |
| Aureococcus anophagefferens |
| Aureoumbra lagunensis |
| Bangiopsis subsimplex |
| Calliarthron tuberculosum |
| Ceramium cimbricum |
| Ceramium japonicum |
| Cerataulina daemon |
| Chaetoceros simplex |
| Chondrus crispus |
| Choreocolax polysiphoniae |
| Coccophora langsdorfii |
| Coeloseira compressa |
| Coscinodiscus radiatus |
| Costaria costata |
| Cryptomonas paramecium |
| Cyanidioschyzon merolae_strain_10D |
| Cyanidium caldarium |
| Cylindrotheca closterium |
| Dasya binghamiae |
| Didymosphenia geminata |
| Durinskia baltica |
| Ectocarpus siliculosus |
| Emiliania huxleyi |
| Erythrotrichia carnea |
| Eunotia naegelii |
| Fucus vesiculosus |
| Galdieria sulphuraria |
| Gelidium elegans |
| Gelidium vagum |
| Gracilaria chilensis |
| Gracilaria chorda |

| |
|---|
| Gracilaria firma |
| Gracilaria tenuistipitata_var._liui |
| Grateloupia taiwanensis |
| Guillardia theta |
| Heterosigma akashiwo |
| Hildenbrandia rivularis |
| Hildenbrandia rubra |
| Kumanoa americana |
| Leptocylindrus danicus |
| Lithodesmium undulatum |
| Mastocarpus papillatus |
| Membranoptera platyphylla |
| Membranoptera tenuis |
| Membranoptera weeksiae |
| Nannochloropsis gaditana |
| Nannochloropsis granulata |
| Nannochloropsis limnetica |
| Nannochloropsis oceanica |
| Nannochloropsis oculata |
| Nannochloropsis salina |
| Nitzschia sp._IriIs04 |
| Odontella sinensis |
| Palmaria palmata |
| Pavlova lutheri |
| Phaeocystis antarctica |
| Phaeocystis globosa |
| Phaeodactylum tricornutum |
| Pleurocladia lacustris |
| Plocamium cartilagineum |
| Porphyridium purpureum |
| Porphyridium sordidum |
| Pseudo-nitzschia multiseries |
| Pyropia haitanensis |
| Pyropia perforata |
| Pyropia pulchra |
| Rhizosolenia imbricata |
| Rhodochaete parvula |
| Rhodomonas salina |
| Rhodymenia pseudopalmata |
| Roundia cardiophora |
| Saccharina japonica |
| Sargassum thunbergii |

| |
|---|
| Schimmelmannia schousboei |
| Schizymenia dubyi |
| Sebdenia flabellata |
| Sporolithon durum |
| Teleaulax amphioxeia |
| Thalassiosira oceanica_CCMP1005 |
| Thalassiosira pseudonana |
| Thalassiosira weissflogii |
| Thorea hispida |
| Toxarium undulatum |
| Trachydiscus minutus |
| Triparma laevis |
| Ulnaria acus |
| Undaria pinnatifida |
| Vaucheria litorea |
| Vertebrata lanosa |
| Wildemania schizophylla |

### Supplementary Table 9. Genomes used to construct the mitochondrial dataset

| Species |
|---|
| Achlya hypogyna |
| Aphanomyces astaci |
| Aphanomyces invadans |
| Asterionella formosa |
| Aurantiochytrium sp._T66 |
| Aureococcus anophagefferens |
| Berkeleya fennica |
| Cafeteria roenbergensis |
| Chattonella marina |
| Chromulina chionophila |
| Colpomenia peregrina |
| Costaria costata |
| Desmarestia viridis |
| Dictyota dichotoma |
| Didymosphenia geminata |
| Ectocarpus siliculosus |
| Fistulifera solaris |
| Florenciella parvula |
| Fragilariopsis cylindrus_CCMP1102 |
| Fucus vesiculosus |
| Heterococcus sp._DN1 |
| Heterosigma akashiwo |
| Laminaria digitata |
| Laminaria hyperborea |
| Monodopsis sp._MarTras21 |
| Nannochloropsis gaditana |
| Nannochloropsis oceanica |
| Navicula ramosissima |
| Ochromonas danica |
| Peronospora tabacina |
| Petalonia fascia |
| Phaeodactylum tricornutum |
| Phaeomonas parva |
| Phytophthora infestans |
| Phytophthora sojae |
| Pleurocladia lacustris |

| |
|---|
| Proteromonas lacertae |
| Pseudo-nitzschia multiseries |
| Pseudopedinella elastica |
| Pseudoperonospora cubensis |
| Pylaiella littoralis |
| Pythium insidiosum |
| Pythium ultimum |
| Saccharina angustata |
| Saccharina japonica |
| Saprolegnia ferax |
| Saprolegnia parasitica_CBS_223.65 |
| Sargassum muticum |
| Sargassum thunbergii |
| Sargassum vachellianum |
| Schizochytrium sp. |
| Scytosiphon lomentaria |
| Skeletonema marinoi |
| Synura peterssenii |
| Synura synuroidea |
| Thalassiosira pseudonana |
| Thraustotheca clavata |
| Trachydiscus minutus |
| Triparma laevis |
| Turbinaria ornata |
| Ulnaria acus |
| Undaria pinnatifida |
| Vischeria sp._CAUP_Q_202 |

### Supplementary Table 10. Genomes used to construct the nuclear dataset

| Species |
|---|
| Albugo candida |
| Aphanomyces astaci |
| Aphanomyces invadans |
| Aplanochytrium kerguelense |
| Aurantiochytrium limacinum |
| Aureococcus anophagefferens |
| Babesia bigemina |
| Babesia bovis |
| Babesia microti_strain_RI |
| Bigelowiella natans |
| Blastocystis hominis |
| Blastocystis sp._NandII |
| Blastocystis sp._subtype_4 |
| Cryptosporidium muris_RN66 |
| Cryptosporidium parvum_Iowa_II |
| Cyclospora cayetanensis |
| Ectocarpus siliculosus |
| Eimeria acervulina |
| Eimeria tenella |
| Fragilariopsis cylindrus |
| Gregarina niphandrodes |
| Hammondia hammondi |
| Ichthyophthirius multifiliis |
| Nannochloropsis gaditana |
| Neospora caninum_Liverpool |
| Oxytricha trifallax |
| Paramecium tetraurelia |
| Perkinsus marinus_ATCC_50983 |
| Phaeodactylum tricornutum_CCAP_1055/1 |
| Phytophthora parasitica |

| |
|---|
| Phytophthora sojae |
| Plasmodiophora brassicae |
| Plasmodium gaboni |
| Plasmodium ovale_curtisi |
| Plasmodium reichenowi |
| Plasmodium vinckei_petteri |
| Plasmodium vivax |
| Plasmopara halstedii |
| Pseudocohnilembus persalinus |
| Pseudo-nitzschia multiseries |
| Reticulomyxa filosa |
| Saprolegnia diclina_VS20 |
| Saprolegnia parasitica_CBS_223.65 |
| Schizochytrium aggregatum |
| Stylonychia lemnae |
| Tetrahymena thermophila_SB210 |
| Thalassiosira oceanica |
| Thalassiosira pseudonana_CCMP1335 |
| Theileria equi_strain_WA |
| Theileria orientalis_strain_Shintoku |
| Theileria parva |
| Toxoplasma gondii_ME49 |
| Vitrella brassicaformis_CCMP3155 |

### Suppl. Figure Legends

#### Suppl. Figure 1 Legend

Complete taxon sampling of the mitochondrial phylogenetic tree inferred using RAxML with LG4X.

#### Suppl. Figure 2 Legend

Complete taxon sampling of the plastid phylogenetic tree inferred using RAxML with LG4X.

#### Suppl. Figure 3 Legend

Complete taxon sampling of the nuclear phylogenetic tree inferred using RAxML with LG4X.

#### Suppl. Figure 4 Legend

Phylogenetic trees of the three compartments based on datasets with common taxonomic sampling and inferred using RAxML with LG4X. A. mitochondrion (22 species); B. plastid (23 species); C. nucleus (23 species).

#### Suppl. Figure 5 Legend

Collapsed trees inferred from supermatrices of 200 nuclear genes selected on the basis of the length of the internal branch leading to E+SSC (RAxML with LG4X model) A. Tree inferred from the supermatrix composed of the 200 genes (39,867 positions) with the shortest E+SSC branch ($SHORT_{nu}$) B. Tree inferred from the supermatrix composed of the 200 genes (47,386 positions) with the longest E+SSC branch ($LONG_{nu}$).

### Suppl. Figure 1

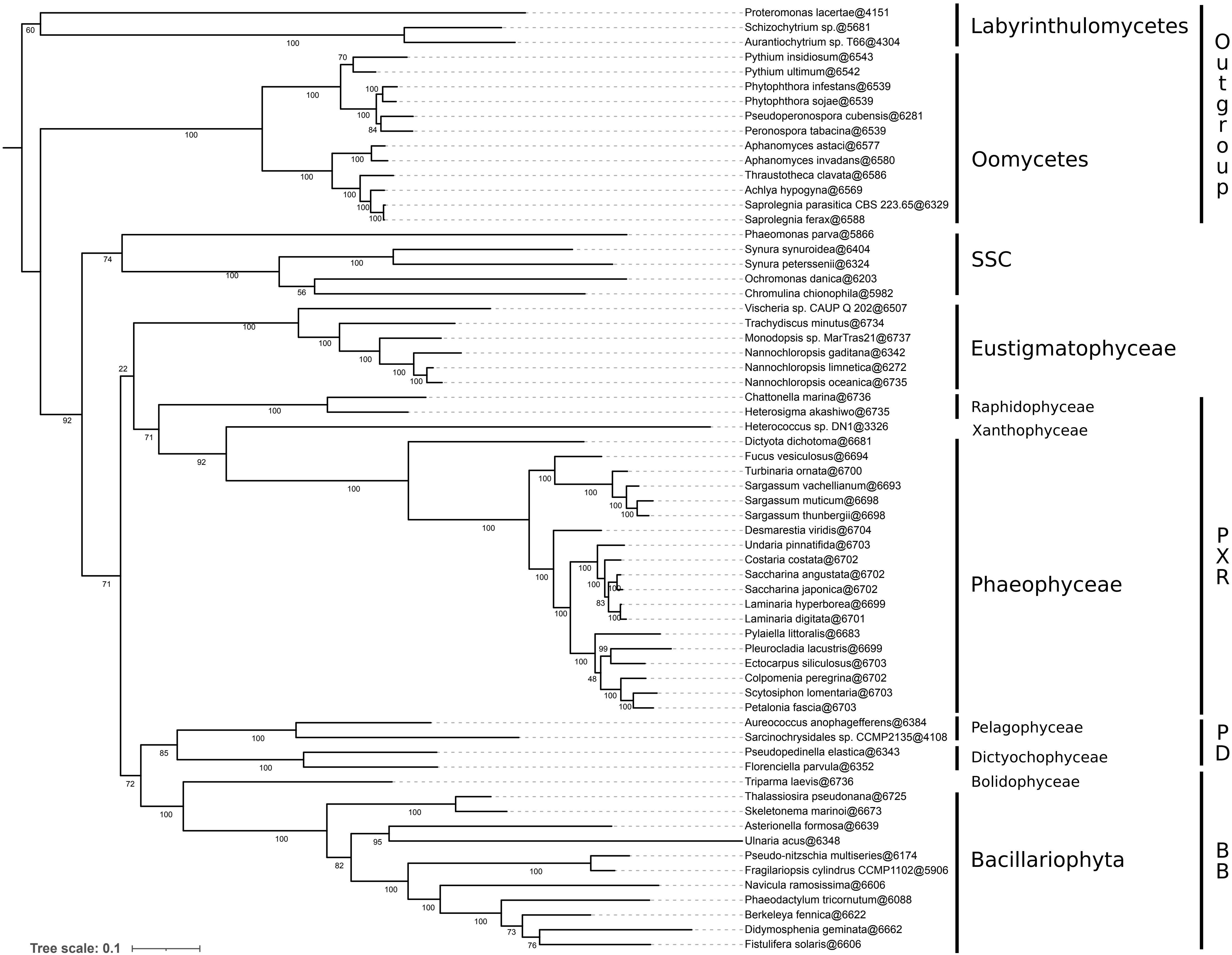

### Suppl. Figure 2

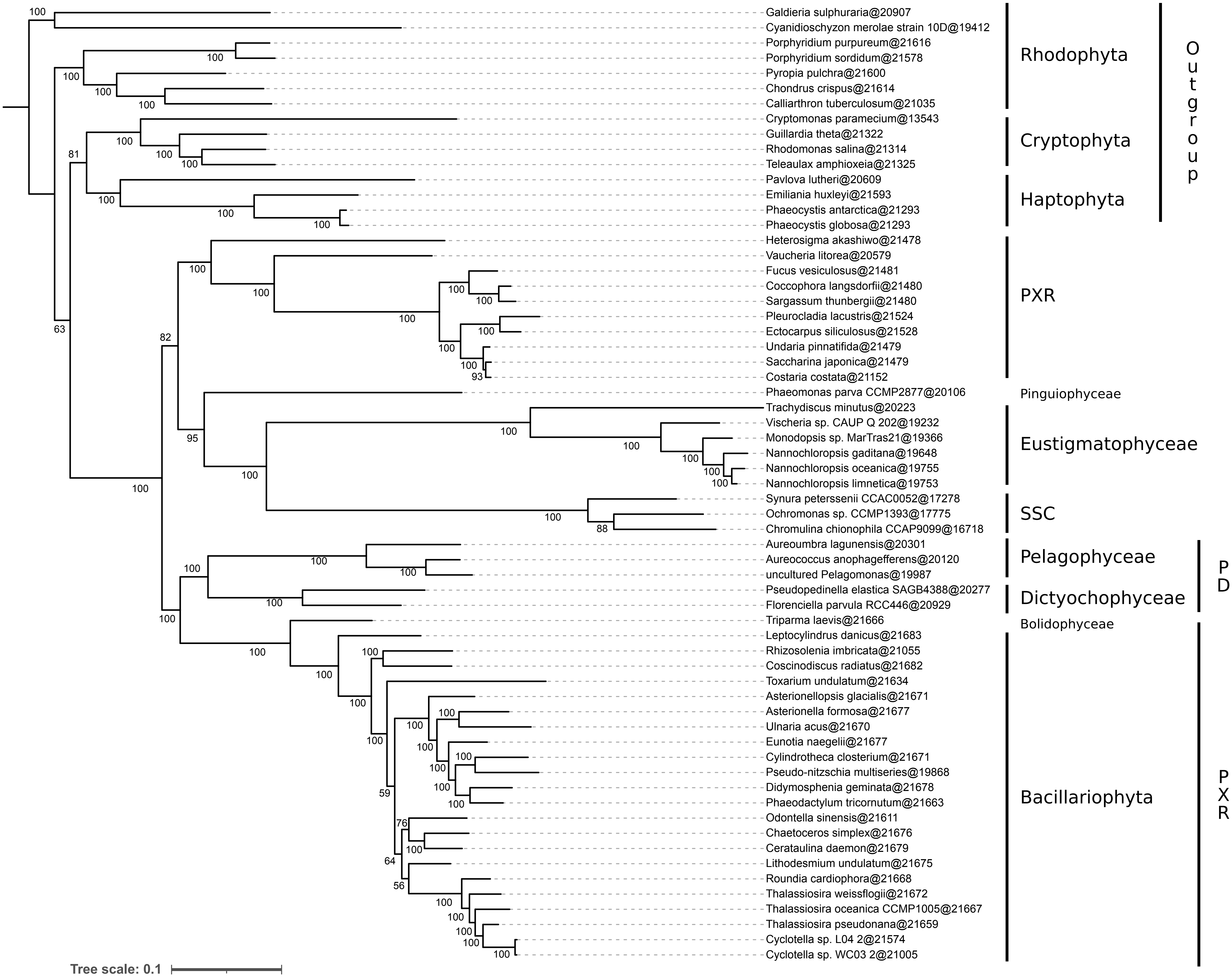

### Suppl. Figure 3

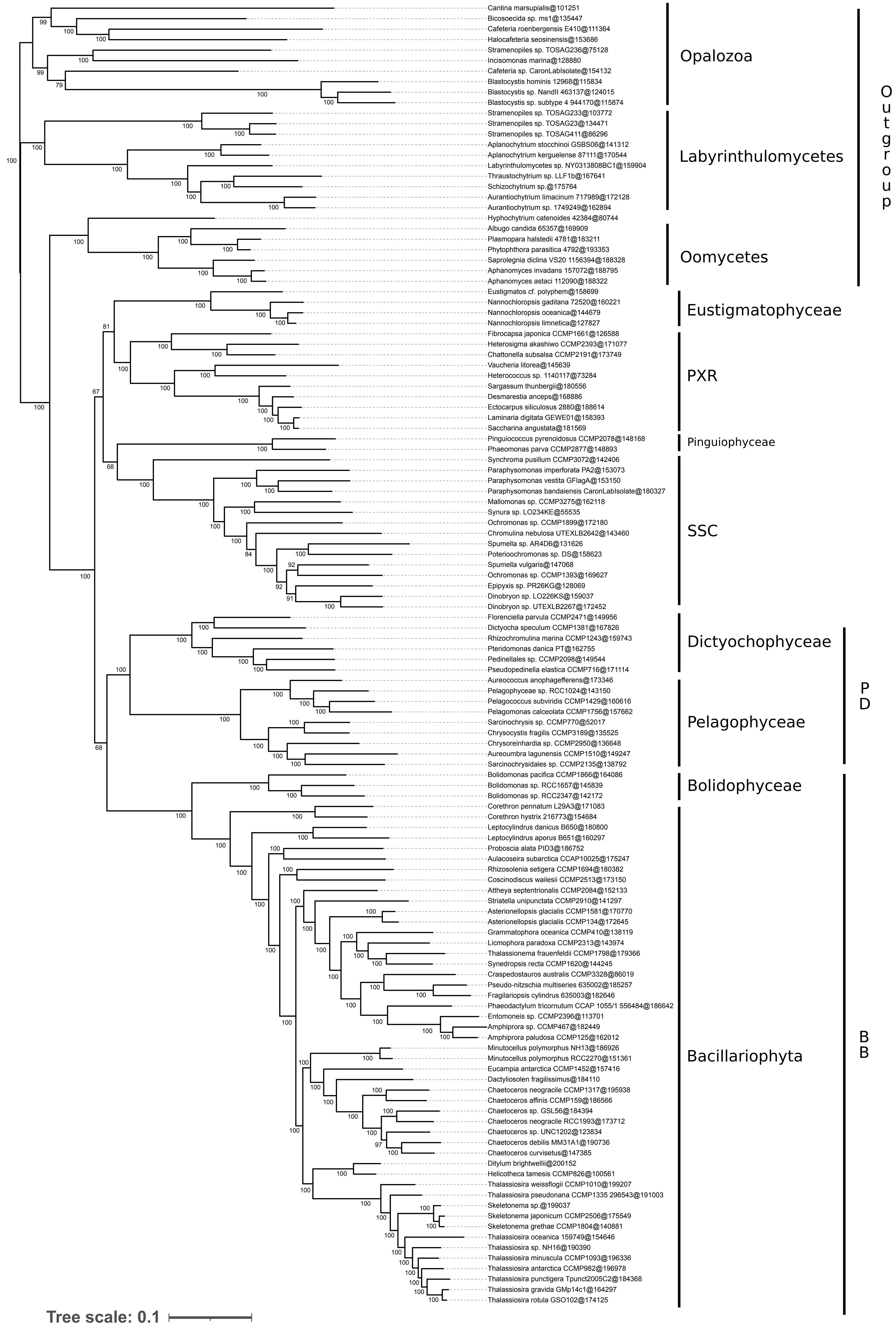

### Suppl. Figure 4

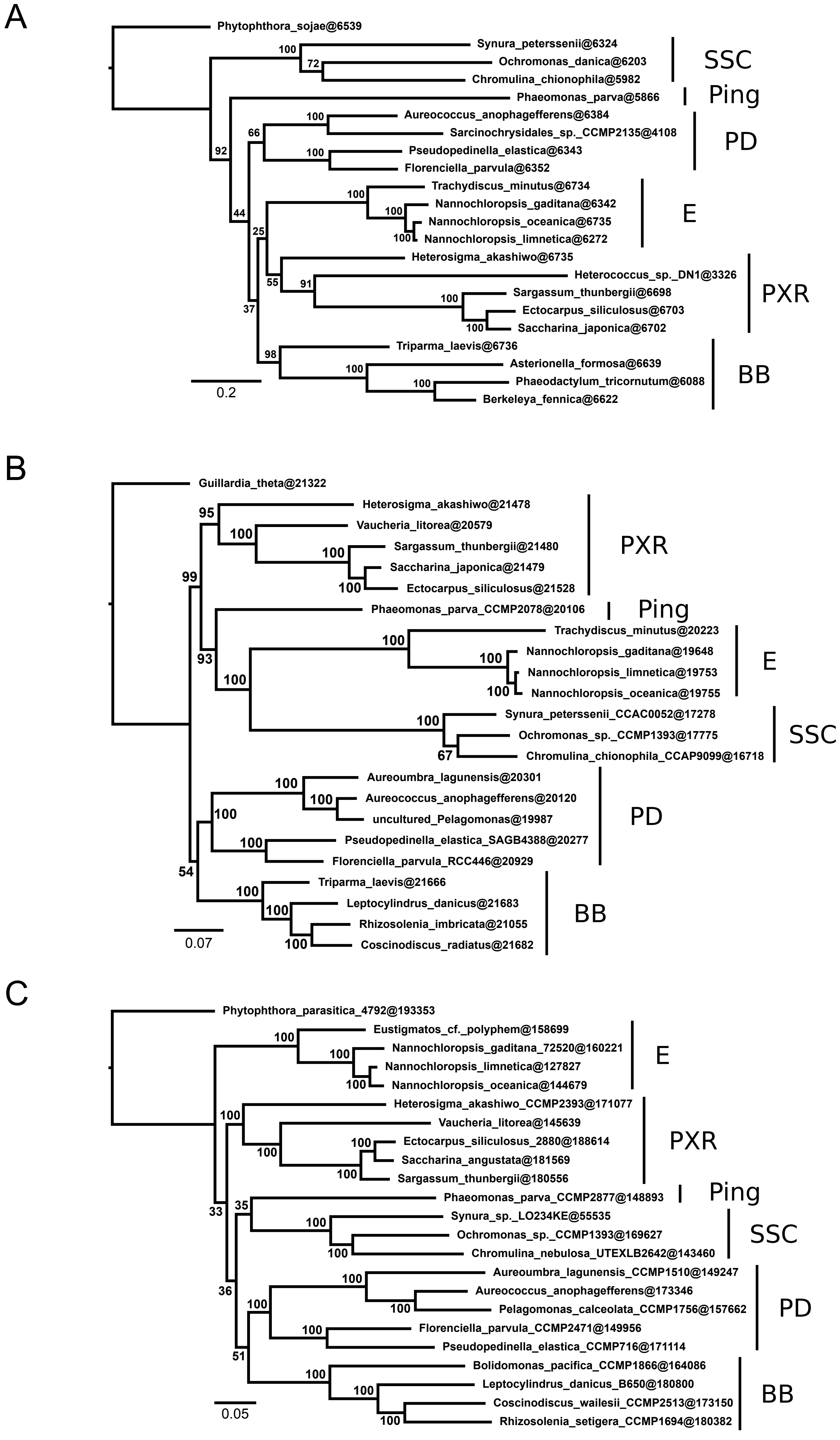

### Suppl. Figure 5

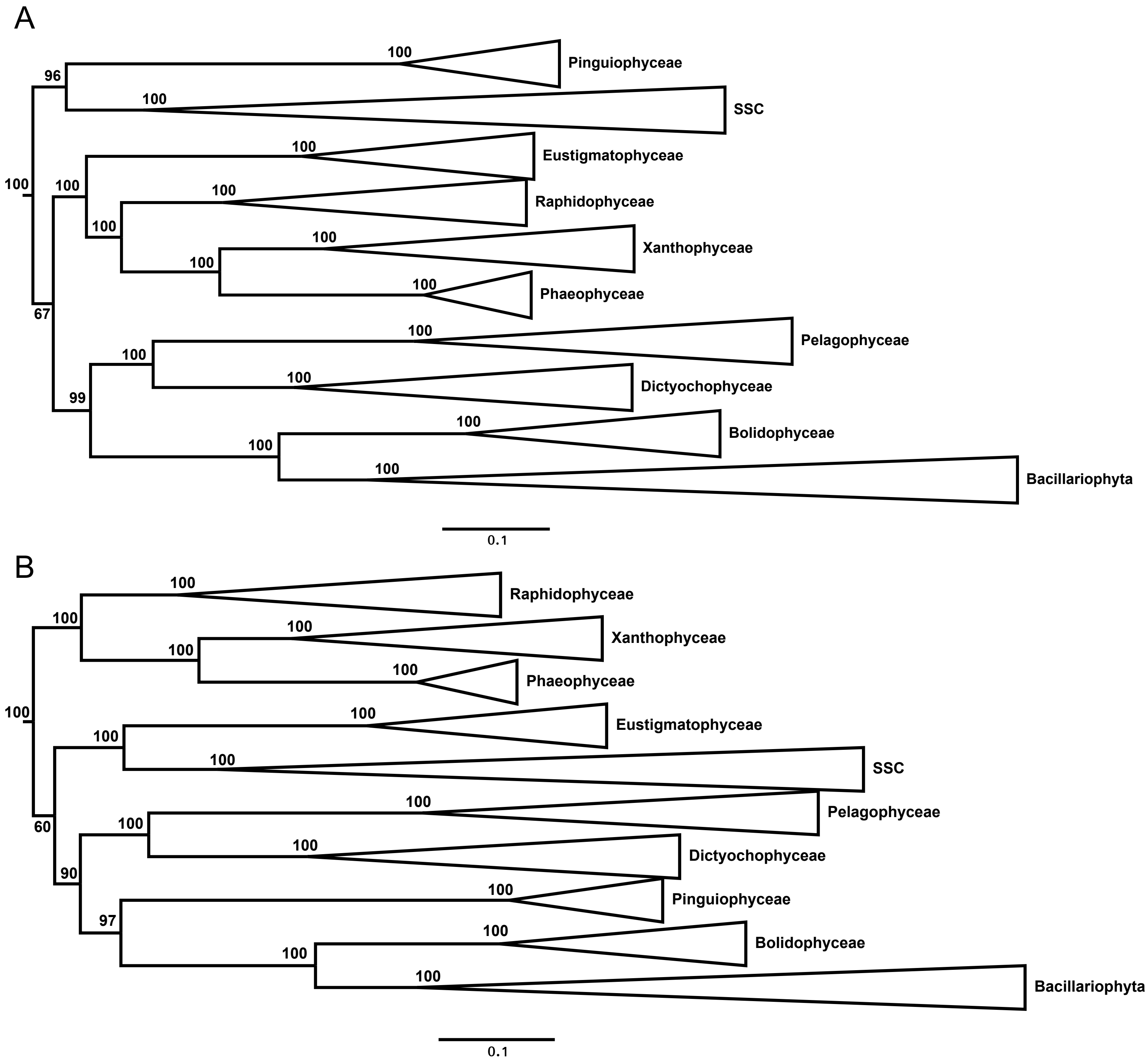
